# Amino Acid Transfer Free Energies Reveal Thermodynamic Driving Forces in Biomolecular Condensate Formation

**DOI:** 10.1101/2024.12.01.625774

**Authors:** Shiv Rekhi, Jeetain Mittal

## Abstract

The self-assembly of intrinsically disordered proteins into biomolecular condensates shows a dependence on the primary sequence of the protein, leading to sequence-dependent phase separation. Methods to investigate this sequence-dependent phase separation rely on effective residue-level interaction potentials that quantify the propensity for the residues to remain in the dilute phase versus the dense phase. The most direct measure of these effective potentials are the distribution coefficients of the different amino acids between the two phases, but due to the lack of availability of these coefficients, proxies, most notably hydropathy, have been used. However, recent work has demonstrated the limitations of the assumption of hydropathy-driven phase separation. In this work, we address this fundamental gap by calculating the transfer free energies associated with transferring each amino acid side chain analog from the dilute phase to the dense phase of a model biomolecular condensate. We uncover an interplay between favorable protein-mediated and unfavorable water-mediated contributions to the overall free energies of transfer. We further uncover an asymmetry between the contributions of positive and negative charges in the driving forces for condensate formation. The results presented in this work provide an explanation for several non-trivial trends observed in the literature and will aid in the interpretation of experiments aimed at elucidating the sequence-dependent driving forces underlying the formation of biomolecular condensates.

## Introduction

Over the past decade, interest in membraneless organelles (MLOs), also known as biomolecular condensates (BCs), has steadily grown due to their potential roles in cellular function^1,2^, neurodegenerative disease^3–6^, and applications as novel biomaterials^7–9^. Several studies have demonstrated the sequence-dependence of MLO formation^10–16^. Coarse-grained (CG) models at the single bead per amino acid resolution have emerged as the primary tool for computational and theoretical investigations of sequence-dependent formation of BCs due to the length and time scales involved in the process of phase separation^17–22^. In these models, sequence-dependence is captured by parameterizing each amino acid using an effective residue-specific interaction potential. The primary challenge in the development of these models is the selection of appropriate effective interaction potentials^21,23,24^.

The most direct measure of these effective potentials are the distribution coefficients or transfer free energies of different amino acids between the dense and the dilute phases (Fig. 1a), but these values are currently unavailable in the literature. An early simplifying assumption to circumvent the lack of these effective potentials was that, similar to the residue-level potentials for protein folding^25–27^, the driving forces for the formation of BCs originate from the hydropathy of the amino acids involved. This led to the adoption of hydrophobicity scales derived from measurements of the free energies of transfer of amino acid side chain analogs from water to vacuum^28^, or from water to a hydrophobic medium such as cyclohexane^29^ as the effective potentials in several sequence-dependent^17,20^ and minimal models for phase separation^30–32^. These models have been extensively used to study the sequence-dependent phase separation of proteins such as the low-complexity (LC) domain of the protein FUS, the RGG domain of LAF1, and the LC domain of hnRNPA1^17–20^. They have also been applied to effects such as the patterning of charged amino acids^12,33,34^ via molecular dynamics (MD) simulations, which led to biophysical insights that were good agreement with experiments.

**Figure 1.**
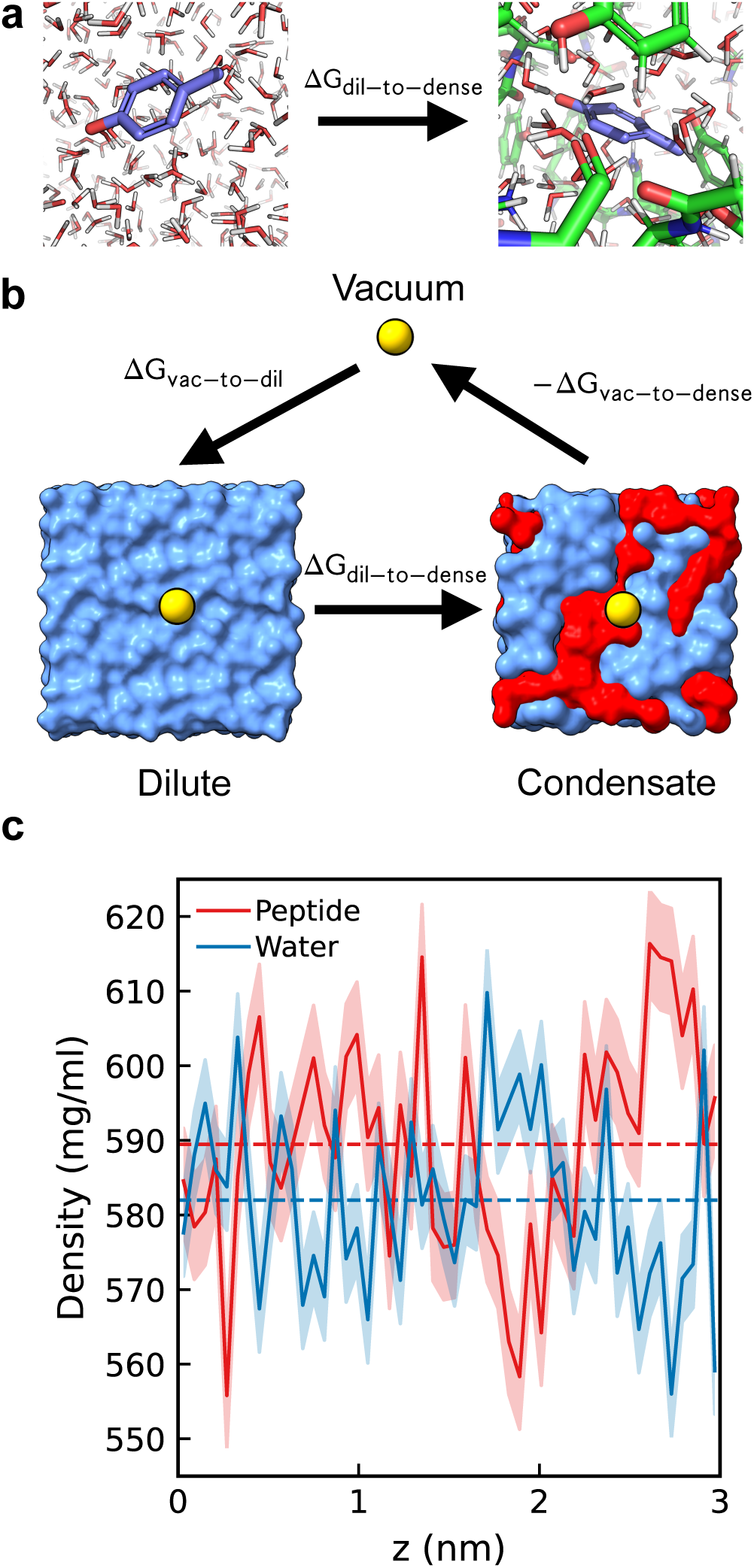
Minimal SYGQ peptides serve as a model system to investigate residue-level driving forces for phase separation. (a) Schematic highlighting the transfer of an amino acid from the dilute to the dense phase (Δ*G_dil_*_−_*_to_*_−_*_dense_*). (b) Schematic of the thermodynamic cycle used to calculate of Δ*G_dil_*_−_*_to_*_−_*_dense_* for all amino acids. (c) Average density profiles of peptide and water along the z dimension of the SYGQ condensate over a 1 simulation. Shaded regions represent block averaged standard error of the mean estimated using 4 200 ns blocks. Dashed lines indicate average peptide and water mass densities within the simulation cell.

Limitations of this simplifying assumption have become apparent as more data from experimental investigations of sequence-dependent phase separation have become available^13,20,23,35^. Solvation free energy, or the transfer free energy of an amino acid from vacuum to the condensed phase, predicts that among the aromatic amino acids, phenylalanine would have the most favorable contribution to phase separation owing to its less favorable solvation compared to tyrosine and tryptophan^28^. However, several investigations have found that tyrosine is significantly favored over phenylalanine^10,12,16^. Also, the difference of ∼60 kJ/mol in experimentally determined solvation free energies of aliphatic and polar amino acid side chain analogs^28^ suggests that aliphatic amino acids would significantly favor phase separation compared to polar residues, but experiments point to a relatively similar contribution of these two classes of amino acids to phase separation^10,35^. To overcome these limitations, residue potentials were optimized using data-driven approaches^20,24,36,37^, pair-potentials between certain residues were augmented to capture interactions such as cation-*π* between cationic and aromatic groups^18,23^, or a different hydrophobicity scale^38^ directly derived from experiments using a host-guest approach was used^19^. However, recent work demonstrates that the limitations of the assumption of hydropathy driven phase separation extend to several other mutations, underlining the need for a robust estimate of residue-level propensities for phase separation^13^.

Here, we address this lack of a fundamental understanding through the calculation of transfer free energies of all amino acid side chains from the dilute phase to the dense phase of a model protein condensate. We find that several non-trivial experimentally observed trends such as the qualitative difference between positive and negatively charged residues^13,39^ and the similar contributions of polar and aliphatic amino acids to phase separation can be explained purely based on the transfer free energies from the dilute phase to the dense phase. Further, we find that the scaffold of the protein condensate can quantitatively alter the transfer free energies but still results in similar rankings of amino acids in terms of their contributions to the driving forces for phase separation. By resolving the contributions resulting from protein- and water-mediated interactions, we uncover a balance between unfavorable hydration and favorable protein-mediated contributions in determining the overall transfer free energies of the individual amino acids. This demonstrates that the tug-of-war between protein and water in condensate formation observed at the chain-level^40^ also exists at the residue-level.

The calculations presented in this work address the fundamental need for a direct measure of residue-level driving forces underlying sequence-dependent phase separation. We believe that the transfer free energies, as well as the methodological approach introduced in this work, represent a significant advance in our understanding of the driving forces underlying phase separation. Further, our findings will aid in the future development of robust physics-based computational models and in the interpretation of experimental results pertaining to the sequence-dependent phase separation of proteins.

## Results and discussion

### Modeling an *in silico* biomolecular condensate

To elucidate residue-level contributions to phase separation, we performed alchemical free energy calculations (see Methods) to quantify the free energy change associated with transferring each amino acid side chain analog from dilute phase into an *in silico* condensate, denoted by Δ*G*_*dil*-*to*-*dense*_ (Fig. 1a). We began by considering only the neutral amino acid side chains. Later, we considered the charged side chains, for which corrections are required to the free energies obtained from alchemical transforms to account for the potential drop across the two phases^41–45^. To compute Δ*G*_*dil*-*to*-*dense*_, we require the transfer from vacuum to the dilute phase, considered as bulk water, Δ*G*_*vac*-*to*-*dil*_, and the transfer from vacuum to a model condensate, Δ*G*_*vac*-*to*-*dense*_(Fig. 1b), such that:

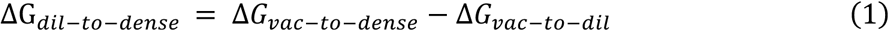

Calculation of Δ*G*_*vac*-*to*-*dil*_, which is equivalent to the solvation free energy has been done for the forcefield and water model used in this work^42,46^ and requires a straightforward simulation procedure (see Methods for details regarding calculation of Δ*G*_*vac*-*to*-*dil*_). However, for the calculation of Δ*G*_*vac*-*to*-*dense*_, an appropriate condensate system is required. Therefore, we first designed a model biomolecular condensate to calculate Δ*G*_*vac*-*to*-*dense*_. A critical consideration was constructing a computationally tractable condensate model that remains representative of real BCs to ensure convergence and relevance of the free energy calculations. We modeled the condensate using a minimal peptide fragment, a common approach in atomistic simulations of BCs^47–49^. Specifically, we selected the tetrapeptide Ser-Tyr-Gly-Gln (SYGQ), capped at the N-terminus with an acetyl group and at the C-terminus with an amine group. This sequence was chosen for its occurrence as a frequent motif in FUS LC and other FET-family proteins^50^, and it captures the polar and aromatic richness characteristic of several intrinsically disordered proteins (IDPs) involved in phase separation, such as hnRNPA1 LC, DDX4 NTD, and TDP43 CTD^50^.

We initialized a cubic simulation box with a side length of 3 nm, containing SYGQ peptides and water molecules in approximately equal mass densities of ∼500 mg/ml, resulting in a total density of ∼1100 mg/ml (Fig. 1c), which is consistent with experimental measurements of FUS LC condensates^51^ and previous all-atom explicit solvent simulations^52^. We conducted 1 *μs* MD simulations in the NPT ensemble to assess the stability and appropriateness of our condensate model. Our simulations revealed a stable condensed phase with minimal fluctuations in peptide and water densities within the simulation box of approximately 50 mg/ml from the mean, indicating the presence of protein-enriched and protein-depleted regions within the condensate (Fig. 1c). This heterogeneity mirrors the complex internal structure of BCs reported in previous atomistic simulations of FUS-LC^52^, and ProTα-H1^53^ as well as recent experiments on minimal *in vitro* reconstituted granular components^54^. Our approach of using SYGQ as a minimal model system, therefore, balances computational efficiency with biological relevance, providing a robust platform for investigating the residue-specific interactions within BCs formed by many IDPs^10–12,16^.

We then performed alchemical free energy calculations to compute the free energy of transferring each of the side chain analogs from vacuum into the model condensate, given by Δ*G*_*vac*-*to*-*dense*_ (see Methods for details regarding alchemical free energy calculations).

### Δ*G*_*vac*-*to*-*dense*_ and Δ*G*_*vac*-*to*-*dil*_ yield contrasting measures of residue-level driving forces

The calculated Δ*G*_*vac*-*to*-*dense*_ values for all neutral amino acid side chain analogs span a range of ∼60 kJ/mol, which is similar to the variations in transfer from vacuum to the dilute phase (Δ*G*_*vac*-*to*-*dil*_). The Δ*G*_*vac*-*to*-*dil*_ values were computed in a solvent box of 3 nm using the same protein force field and simulation parameters as Δ*G_*vac*-*to*-*dense*_*, and they align with experiments quantifying the solvation free energy of amino acid side chain analogs^28^ (Fig. 2a, Supplementary Fig. S1). We observed that certain configurations for some amino acids yielded Δ*G_*vac*-*to*-*dense*_* values outside of 1.5 times the interquartile range of all configurations, highlighting the extent to which Δ*G_*vac*-*to*-*dense*_* can vary based on the local environment within the condensate.

**Figure 2.**
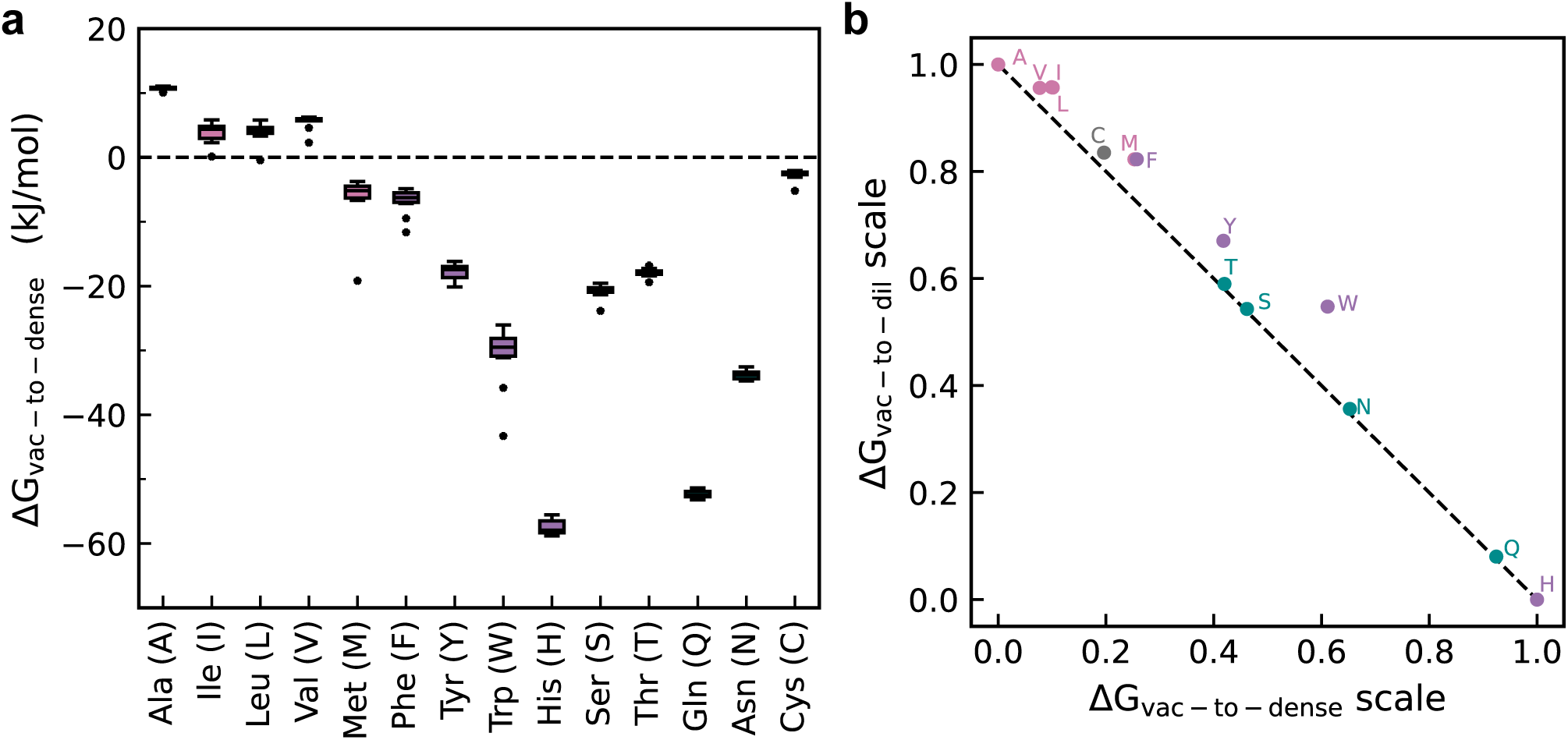
Transfer from vacuum to condensate and transfer from vacuum to solvent are correlated. (a) Transfer from vacuum to condensate (Δ*G_vac_*_−_*_to_*_−_*_dense_*) of all neutral amino acids in kJ/mol. Amino acids are grouped by side chain properties: aliphatic, aromatic, polar, and cysteine. Horizontal lines in the box plots represent the median values of 10 different insertions; whiskers extend to 1.5 times the interquartile region. Data points falling outside the whiskers are indicated by asterisks. (b) Comparison of Δ*G_vac_*_−_*_to_*_−_*_dense_* and transfer from vacuum to solvent (Δ*G_vac_*_−_*_to_*_−_*_dil_*) for neutral amino acid side chains. Values are normalized from 0-1 such that 0 indicates the least favorable contribution to phase separation while 1 indicates the most favorable. Amino acids are labeled with their corresponding single-letter amino acid code.

Our results show that only the aliphatic amino acids alanine, isoleucine, leucine, and valine have unfavorable (positive) Δ*G_*vac*-*to*-*dense*_*, with alanine being the most unfavorable. In contrast, histidine exhibits the most favorable Δ*G_*vac*-*to*-*dense*_* (Fig. 2a). Among the remaining aromatic amino acids, tryptophan is the most favorable, followed by tyrosine and phenylalanine. Polar amino acids also display highly favorable Δ*G_*vac*-*to*-*dense*_* values, with glutamine and asparagine being significantly more favorable than serine and threonine. Notably, methionine has a significantly more favorable Δ*G_*vac*-*to*-*dense*_* compared to other aliphatic amino acids, possibly due to the presence of a sulfur atom in its side chain.

Direct comparison between Δ*G_*vac*-*to*-*dense*_* and Δ*G_*vac*-*to*-*dil*_* surprisingly reveals that Δ*G_*vac*-*to*-*dense*_* scales with Δ*G_*vac*-*to*-*dil*_* resulting in a very high quantitative correlation between the two (Supplementary Fig. S1). Interestingly, considering the context of driving forces for phase separation, more negative Δ*G_*vac*-*to*-*dil*_* indicates reduced contributions to phase separation, while more negative Δ*G_*vac*-*to*-*dense*_* indicates enhanced contributions to phase separation. We normalize Δ*G_*vac*-*to*-*dil*_* and Δ*G_*vac*-*to*-*dense*_* values between 0 and 1, where 0 represents the least favorable and 1 represents the most favorable contribution to phase separation. This normalization revealed that Δ*G_*vac*-*to*-*dil*_* and Δ*G_*vac*-*to*-*dense*_* predict opposite residue-level contributions to phase separation (Fig. 2b). We see that the residues predicted to be most favored for phase separation from Δ*G_*vac*-*to*-*dil*_* (aliphatic) are the most disfavored from Δ*G_*vac*-*to*-*dense*_* (Fig. 2b). More notably, a number of the shortcomings of Δ*G_*vac*-*to*-*dil*_* remain in Δ*G_*vac*-*to*-*dense*_*. Namely, the large difference between polar and aliphatic amino acids, as well as the prediction of aromatic residues lying in the middle of the scale, contrary to their observed importance in phase separation^10,35,55^, highlights that Δ*G_*vac*-*to*-*dense*_*, like Δ*G_*vac*-*to*-*dil*_*, cannot adequately describe residue-level contributions to phase separation thermodynamics (Fig. 2b). This is most likely due to the use of vacuum as the reference state in calculating Δ*G_*vac*-*to*-*dense*_*, whereas the calculation of Δ*G_*dil*-*to*-*dense*_* considers the reference state as the dilute phase. Therefore, we next considered whether calculating the transfer of each amino acid side chain from bulk water dilute phase to the condensate would yield a better description of the driving forces underlying the formation of condensates.

### Transfer free energy from bulk water to condensate (Δ*G_*dil*-*to*-*dense*_*) captures the thermodynamics of phase separation

We calculated the transfer free energy of all amino acid side chains from bulk water to the condensate, denoted as Δ*G_*dil*-*to*-*dense*_*, using a thermodynamic cycle (Fig. 1b) as given by equation 1. The calculated Δ*G_*dil*-*to*-*dense*_* values for all neutral amino acid side chains show a variation of approximately 20 kJ/mol between the most unfavorable and most favorable amino acids (Fig. 3a). Similar to the Δ*G_*vac*-*to*-*dense*_* results, alanine exhibits the most unfavorable Δ*G_*dil*-*to*-*dense*_*; however, unlike in the Δ*G_*vac*-*to*-*dense*_* results, tryptophan shows the most favorable Δ*G_*dil*-*to*-*dense*_* value instead of histidine (Fig. 3a). Notably, alanine is the only one with an unfavorable Δ*G_*dil*-*to*-*dense*_*, supporting frequent experimental observations of reduced phase separation upon mutations of diverse residue types to alanine^13,56,57^. This is except in cases where secondary-structure-mediated interactions may contribute to phase separation (e.g. helix-helix interactions in TDP43 CTD^4,58^).

**Figure 3.**
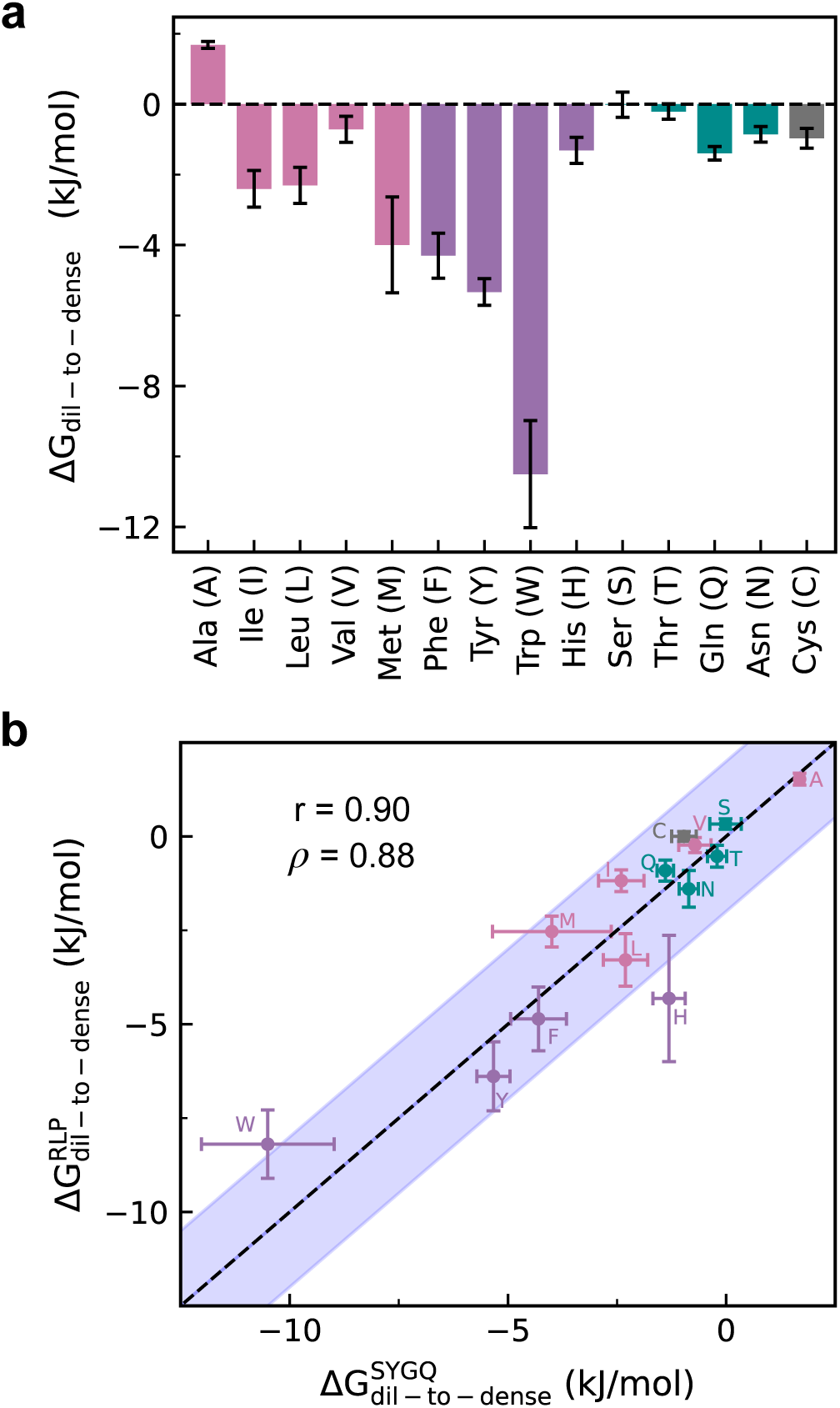
Δ*G_dil−to−dense_* quantifies residue-level driving forces in accordance with experimental knowledge. (a) Transfer free energy from the dilute to the dense phase (Δ*G_dil_*_−_*_to_*_−_*_dense_*) for all neutral amino acid side chains. Amino acids are grouped as in Fig. 2a. Error bars represent the standard error of mean (SEM) of Δ*G_dil_*_−_*_to_*_−_*_dense_* calculated over 10 unique insertion positions. (b) Comparison of Δ*G_dil_*_−_*_to_*_−_*_dense_* calculated from the condensates composed of the SYGQ (x axis) and RLP (y axis) repeat unit peptides. Shaded areas denote +/- 2 kJ/mol from the diagonal. Errors for both condensate systems are reported as the SEM of 10 different insertion positions. Pearson (r) and Spearman correlation coefficients (ρ) are shown as measures of correlation.

The calculated Δ*G_*dil*-*to*-*dense*_* values mirror several other experimental trends reported in the literature, extending beyond the unfavorability of alanine in the context of phase separation. We observe that polar and aliphatic amino acids are now separated by a range of approximately 4 kJ/mol – more than an order of magnitude less than the 60 kJ/mol difference observed in Δ*G_*vac*-*to*-*dense*_* (Fig. 2a, Fig. 3a). This relatively similar contribution of polar and aliphatic amino acids aligns with observations of their comparable effects on phase separation, as seen in studies on FUS LC, hnRNPA1 LC, and polar-rich artificial IDPs^10,13,15,16^. Among the aliphatic amino acids, methionine has the most favorable Δ*G_*dil*-*to*-*dense*_* (Fig. 3a), consistent with the observations for TDP43 CTD, in which the methionine residues were found to engage in a significant number of protein-protein contacts, playing a key role in its phase separation^57^.

Among the polar residues, glutamine is the most favorable (Fig. 3a), supporting recent findings regarding its importance compared to other polar residues in promoting the condensate formation of FUS LC^59^. Our results also reveal that similar Δ*G_*dil*-*to*-*dense*_* values for different amino acids can arise from different mechanisms. For aliphatic amino acids (except methionine), unfavorable Δ*G_*vac*-*to*-*dense*_* and Δ*G_*vac*-*to*-*dil*_* lead to a favorable Δ*G_*dil*-*to*-*dense*_* due to their relative magnitudes (Supplementary Fig. S2). In other words, the condensate environment is less unfavorable for these residues than bulk water. In contrast, for polar amino acids, both Δ*G_*vac*-*to*-*dense*_* and Δ*G_*vac*-*to*-*dil*_* are highly favorable, with Δ*G_*vac*-*to*-*dense*_* being slightly more favorable than Δ*G_*vac*-*to*-*dil*_*, resulting in Δ*G_*dil*-*to*-*dense*_* values comparable to those of the aliphatic amino acids (Supplementary Fig. S2).

The most favorable Δ*G_*dil*-*to*-*dense*_* among the neutral amino acids are observed for the aromatic residues (WYF) in the order tryptophan >> tyrosine > phenylalanine (Fig. 3a), consistent with the known prominent role of these residues in phase separation and their relative ordering^12,13,60^. In contrast, histidine, despite being aromatic, has a significantly less favorable Δ*G_*dil*-*to*-*dense*_*; we find that favorable Δ*G_*vac*-*to*-*dense*_* is almost completely offset by highly favorable Δ*G_*vac*-*to*-*dil*_*, similar to polar residues (Supplementary Fig. S2, Fig. 3a). Although Δ*G_*vac*-*to*-*dil*_* is favorable for phenylalanine, tyrosine, and tryptophan, Δ*G_*vac*-*to*-*dense*_* is significantly more favorable, leading to favorable Δ*G_*dil*-*to*-*dense*_*^12,55,60^ (Supplementary Fig. S2, Fig. 3a). However, to gain mechanistic insight into the differences between these residues in the context of driving forces for phase separation, it is necessary to separate the protein- and water-mediated contributions to their transfer free energies, as we do later in this article.

### *ΔG_*dil*-*to*-*dense*_* remains relatively unchanged for a different model condensate

Given the expected role of protein-mediated interactions in determining Δ*G_*vac*-*to*-*dense*_*, we investigated how changing the protein sequence of the condensate affects the values of Δ*G_*dil*-*to*-*dense*_*. We used the resilin-like polypeptide (RLP) repeat unit^15^, with the sequence GRGDSPYS, capped at the N- and C-termini. The RLP repeat unit retains the polar-rich nature of the SYGQ sequence while introducing a pair of oppositely charged amino acids. Like the SYGQ sequence, the RLP repeat unit captures compositional biases prevalent in proteins that form BCs^13^. We observed that the Δ*G_*dil*-*to*-*dense*_* values between the two condensates are very similar, with most amino acids lying on the diagonal line within statistical uncertainty and all lying within a range of ±2 kJ/mol (Fig. 3b).

The comparison of Δ*G_*dil*-*to*-*dense*_* between the SYGQ and RLP condensates highlights that, in both condensates, aromatic amino acids remain the most favorable, polar and aliphatic amino acids have similar transfer free energies, and alanine remains unfavorable. This suggests that a dominant factor in determining Δ*G_*dil*-*to*-*dense*_* is the difference in microenvironments between the dilute and dense phases. This is also in line with recent work investigating the partitioning of small molecules into condensates^61,62^. However, the transfer free energies from the dilute phase to the dense phase between the two condensate systems are not equivalent for all amino acids (Fig. 3b). For instance, within the SYGQ condensate, asparagine has a less favorable transfer than glutamine, while isoleucine and leucine show similar transfer free energies (Fig. 3a, 3b). However, within the RLP condensate, we see that asparagine is slightly more favorable than glutamine, and leucine is more favorable than isoleucine (Fig. 3b). This demonstrates that despite both peptide sequences having similar polar-rich character, differences in the amino acid sequence of the condensate can alter residue-level driving forces for phase separation. This can lead to context-dependent effects, such as the opposite effects of glutamine-to-asparagine mutations observed in FUS LC^59^ and hnRNPA1 LC^16^.

### *ΔG_*dil*-*to*-*dense*_* correlates with experimental and data-derived hydropathy scales

MD simulations based on CG models have become indispensable for studying sequence-dependent protein phase separation. An early starting point in the development of CG models for phase separation involved the use of the Kim-Hummer (KH) potential^63^, which is based on the Miyazawa-Jernigan (MJ) contact potential^27^. The MJ potential is derived from contact statistics of protein structures within the PDB database. Consistent with many previous studies^23,64^ that have highlighted the discrepancies associated with using MJ-based models to study phase separation, we find that the MJ potential values for self-interaction of neutral amino acids do not correlate well with our calculated transfer free energies (Fig. 4a). This indicates that energetics derived to describe the thermodynamics of protein folding may not be transferable to phase separation^23,64^. To further investigate, we compared our transfer free energies to two state-of-the-art CG models developed for phase separation: HPS-Urry^19,38^ and CALVADOS2^24^. We find that the transfer free energies from the dilute phase to the dense phase correlate strongly with both CG model scales, and the ranking of amino acids is in good agreement (Figs. 4b and 4c). This correlation with experimentally derived (HPS-Urry) and data-derived (CALVADOS2) residue-level interaction scales reinforces the conclusion that our fundamental approach of calculating transfer free energies from the dilute phase to the dense phase effectively uncovers the residue-level driving forces for phase separation without the requirement of limited experimental data as inputs.

**Figure 4.**
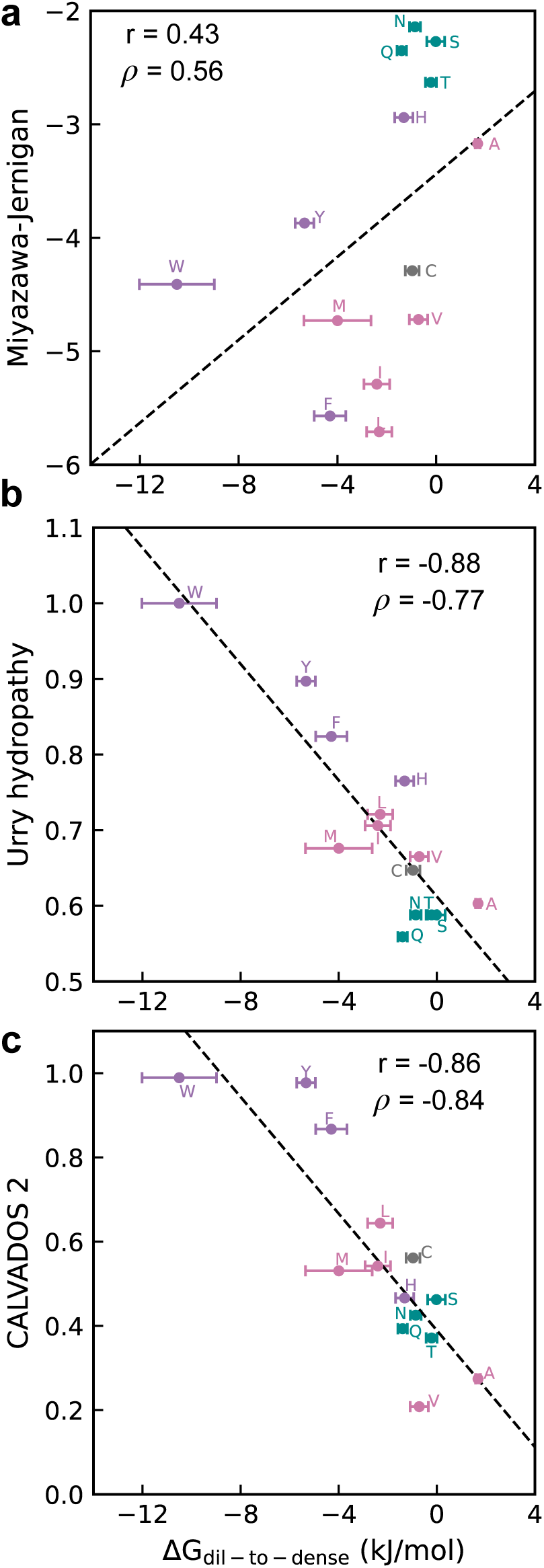
Δ*G_dil−to−dense_* correlates with existing phase separation scales. Correlation of transfer free energy from the dilute to the dense phase (Δ*G_dil_*_−_*_to_*_−_*_dense_*) of all neutral amino acid side chain analogs with (a) Miyazawa-Jernigan self-interaction parameters, (b) Urry hydropathy values, and (c) CALVADOS2 hydropathy values. Pearson (r) and Spearman (ρ) correlation coefficients are shown as a measure of correlation, while the dashed lines indicate the lines of best fit.

### Separation of Δ*G_*dil*-*to*-*dense*_* into contributions from protein-mediated and water-mediated interactions

The microenvironment inside the condensate consists of both protein and water molecules, which collectively dictate the overall transfer free energies of the amino acids. To further understand these interactions within the condensate, we assume that the net transfer free energy from the dilute phase to the dense phase of the condensate is the result of additive contributions of protein- and water-mediated interactions within the condensate.

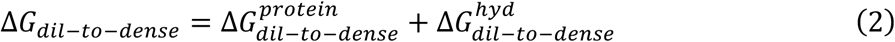

We then calculated the transfer free energy from the bulk water to a condensed phase composed purely of protein (“dry” condensate), with all water molecules removed, denoted as 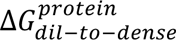 (details of system preparation and equilibration are provided in the Methods section) using a similar thermodynamic cycle-based approach (Fig. 5a). The aromatic amino acids, phenylalanine, tryptophan, and tyrosine, show the most favorable transfer from the dilute phase into the dry condensate, consistent with the high protein contact propensities of these amino acids observed in atomistic simulations of biomolecular condensates^13,47,52,59^. Interestingly, we observe a good qualitative correlation between Δ*G_*dil*-*to*-*dense*_* and 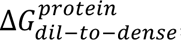, although 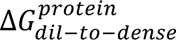 is significantly more favorable for all amino acids, including alanine (Fig. 5b).

**Figure 5.**
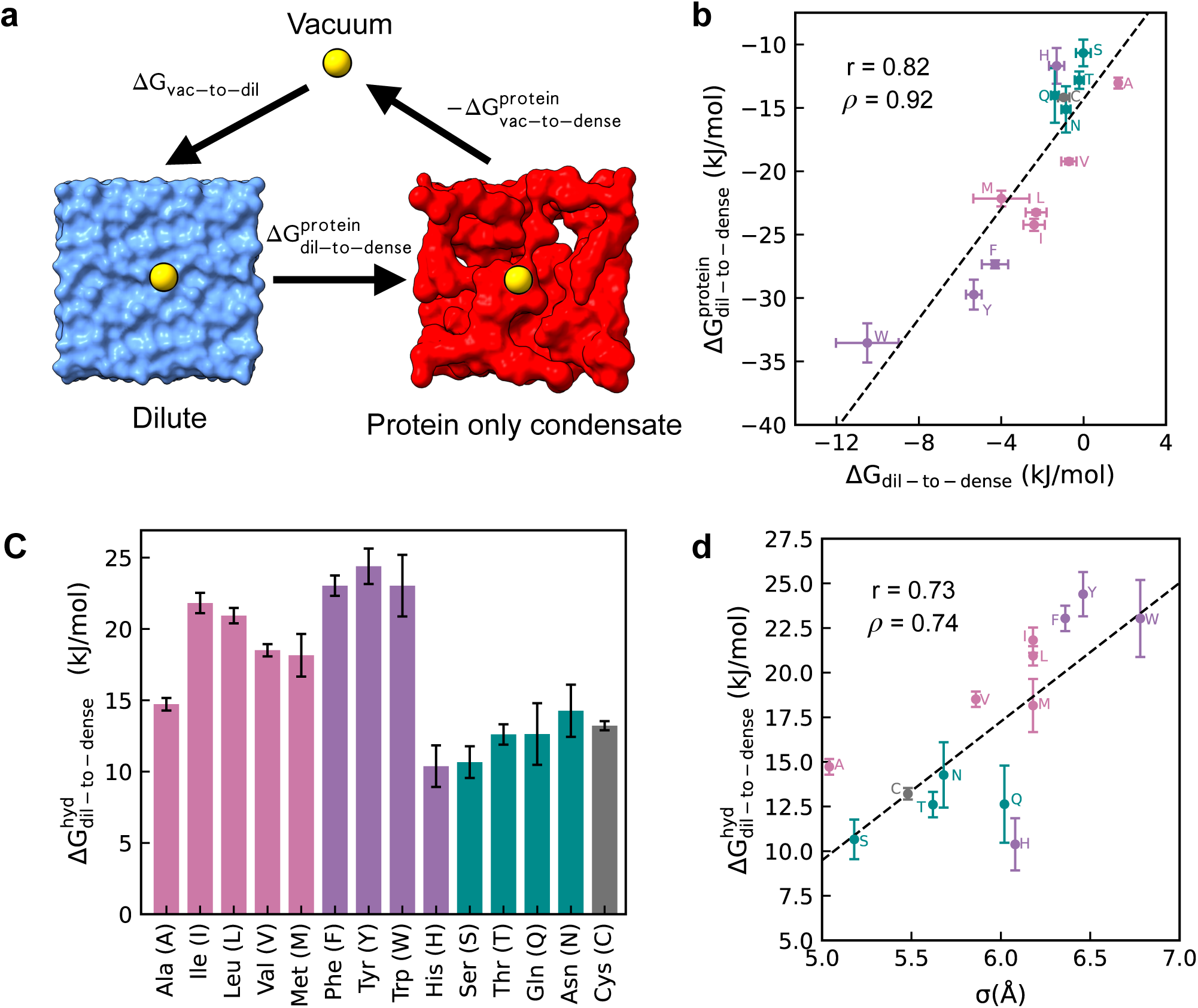
Transfer free energies depend on both protein and water molecules. (a) Schematic highlighting the thermodynamic cycle used for the calculation of the protein mediated contribution to transfer free energy from the dilute to the dense phase 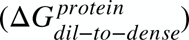 (b) Correlation between transfer free energy from the dilute to the dense phase (Δ*G_dil_*_−_*_to_*_−_*_dense_*) and 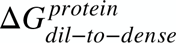 for all neutral amino acid side chains. Error bars for 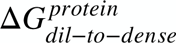 are calculated as SEM over 5 configurations, whereas for Δ*G_dil_*_−_*_to_*_−_*_dense_*, they are calculated over 10 configurations. Pearson (r) and Spearman (ρ) correlation coefficients are shown as measures of correlation. Dashed line shows the line of best fit. (c) Hydration mediated contribution to transfer free energy from the dilute to the dense phase 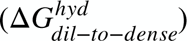 in kJ/mol for all neutral amino acids. (d) Δ*G_dil_*_−_*_to_*_−_*_dense_* plotted against amino acid diameters (σ) in Å for all neutral amino acids. Pearson (r) and Spearman (ρ) correlation coefficients are shown as measures of correlation. Dashed black line shows line of best fit.

The difference between the transfer from the bulk water phase into the condensate and the dry condensate, 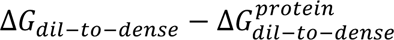, can be used to estimate the role of water-mediated interactions within the condensate to the transfer free energy, referred to as the hydration part of the transfer free energy 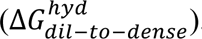. We found that 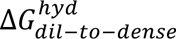, which represents transfer from bulk water to water inside the condensate, is highly unfavorable^40^ for all amino acids (Fig. 5c). Furthermore, we find that the transfer from vacuum to the bulk water, Δ*G_*vac*-*to*-*dil*_*, does not qualitatively explain the relative magnitudes of 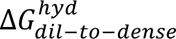 for the different amino acids (Supplementary Fig. S3) suggesting that water within the condensed phase is distinct from bulk water. We characterize the tetrahedrality^65^ of water within the condensed phase and find that the hydrogen bonding network of water within the condensed phase is different from bulk water (Supplementary Fig. S3), due to confinement and direct interactions between water and protein^40^. We reason that a consequence of this confinement is reduced density fluctuations, leading to lower probabilities of creating a solute-sized cavity, resulting in unfavorable 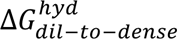 for all neutral amino acids^66–68^. We observe that the magnitude of 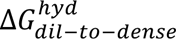 increases as the amino acid gets larger reflecting the entropic cost of cavity formation for larger solutes^69,70^ (Fig. 5d).

Using the values of 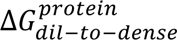 and 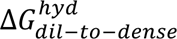, we revisit the ranking of tryptophan, tyrosine and phenylalanine in their favorability of transfer from the dilute to the dense phase. Recent work has suggested that the preference for tyrosine over phenylalanine in condensate formation is primarily driven by the higher hydration within the condensate microenvironment^64^. Considering the comparison between phenylalanine and tyrosine, we see that the water-mediated 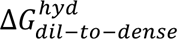 is more unfavorable for tyrosine, consistent with its larger size, (Fig. 5c,5d), but the protein-mediated contribution from 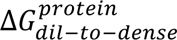 is more favorable, consistent with the higher self- and cross-interactions of tyrosine vs. phenylalanine^18^ (Fig. 5b).

Therefore, our analysis suggests that protein-mediated interactions within the polar rich peptide condensate environment lead to the preference of tyrosine over phenylalanine in the context of phase separation (Fig. 3a). We next investigate why tryptophan shows significantly more favorable transfer into the condensate as compared to tyrosine despite more favorable solvation in the computational model. We observe that the 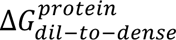 contribution for tryptophan is ∼5 kJ/mol more favorable than for tyrosine (Fig. 5b) but the 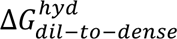 is lower despite an increase in size of ∼0.25 Å (Fig. 5c, 5d). The mechanistic picture that emerges is that both tryptophan and tyrosine form significantly more contacts, however the favorable protein-mediated contribution is offset to a lower extent by the unfavorable water-mediated contribution for tryptophan compared to tyrosine leading to an enhanced favorability of transfer from the dilute phase to the dense phase of the condensate (Fig. 3a).

Our analysis of the neutral amino acids reveals an interplay between bulk-like water within the dilute phase, protein within the condensed phase, as well as confined water within the condensed phase collectively determine the transfer from the dilute phase to the condensed phase.

### Transfer free energy of charged amino acid side chain analogs uncovers charge asymmetry

For charged amino acids, the free energies measured using alchemical transforms such as those done in this work require a correction relating to the energetic cost of crossing the interface between the two media, originating from their different electrostatic potentials. Namely, intrinsic free energies estimated from alchemical free energy calculations are corrected by a surface potential term,

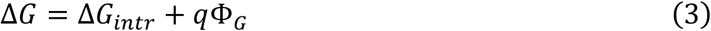

Where, q is the charge of the species and Φ_*G*_ is the Galvani potential jump between the two phases, given by *Fχ* where F is Faraday’s constant and *χ* is the electrostatic potential difference between the two phases or the phase potential^41,42^. To account for this, we computed the electrostatic potentials of both the dilute phase (*χ*^*dil*^) and the dense phase (*χ*^*dense*^) with respect to the reference vacuum state, where the electrostatic potential is set to 0 (see Methods for more details). We find that the Galvani potentials for the dilute phase 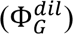 and dense phase 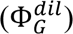 are −48.06 and −27.55 kJ/mol/e.

The Δ*G_*dil*-*to*-*dense*_* values for charged amino acids reveal a distinct difference based on the sign of the charge (Fig. 6a) with positively charged amino acids strongly favoring partitioning into the condensate. Δ*G_*vac*-*to*-*dil*_* and Δ*G_*vac*-*to*-*dense*_* are both highly favorable for the charged amino acids (Supplementary Fig. S4), but in the case of negatively charged amino acids, the highly favorable solvation^71^ overcomes Δ*G_*vac*-*to*-*dense*_*, leading to unfavorable Δ*G_*dil*-*to*-*dense*_*. This charge asymmetry observed in Δ*G_*dil*-*to*-*dense*_* provides a compelling explanation for several experimental observations. Recent studies have demonstrated that a positively charged A-IDP can undergo phase separation whereas a negatively charged A-IDP cannot under the same experimental conditions, despite having the same charge magnitude^13,14^. Notably, this behavior for polycationic peptides persists even up to a net charge per residue of 0.3^72^. Furthermore, the disordered region of the protein MED1, which plays a critical role in transcriptional condensates, contains a significant fraction of positively charged amino acids and undergoes homotypic phase separation *in vitro*^56^. Similarly, the N-terminal domain of the protein DDX4, which at neutral pH carries a net negative charge, demonstrates the highest propensity to phase separate at a charge state of +13^73^. Additionally, two engineered variants of green fluorescent protein (GFP) with net positive and net negative charges were found to exhibit differential partitioning into condensates, a phenomenon attributed to the promiscuity of interactions involving positively charged amino acids^39^. More broadly, analysis of LLPSDB, a database dedicated to proteins undergoing liquid-liquid phase separation, revealed that sequences capable of phase separation are enriched in cationic amino acids, while those that do not phase separate are enriched in anionic amino acids^74^. The above findings have been attributed to the ability of positively charged amino acids to participate in cation-*π* interactions with aromatic groups, or complex coacervation of protein and negatively charged RNA or DNA molecules within cellular condensates. However, here we show that the widespread preference for positive charge within condensate-forming proteins may also originate from their preference for the dense phase over the dilute phase.

**Figure 6.**
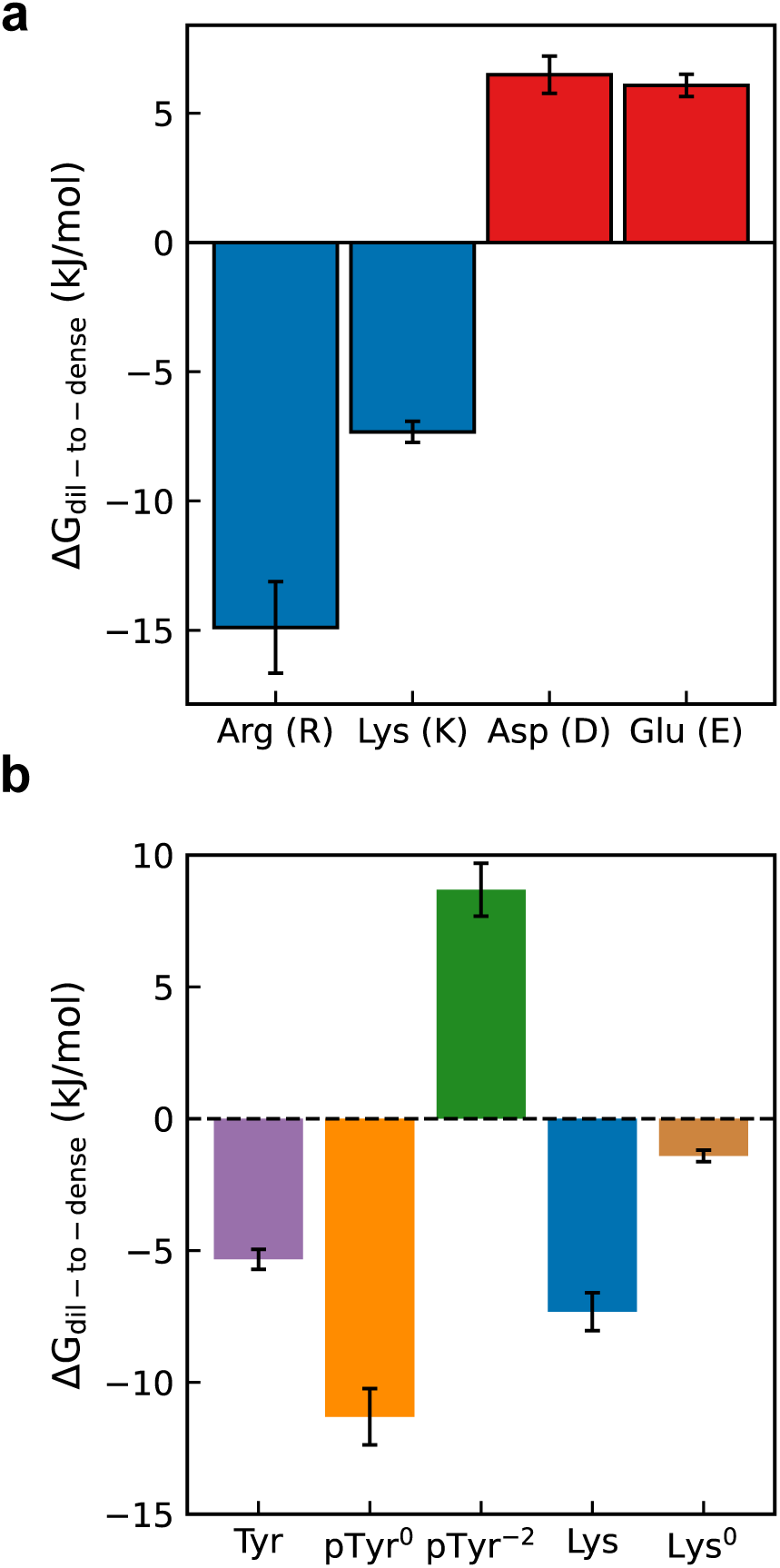
Positive and negatively charged amino acids show differential partitioning into the condensed phase. (a) Transfer free energy from the dilute to the dense phase of the SYGQ condensate (Δ*G_dil_*_−_*_to_*_−_*_dense_*) for the charged amino acids in kJ/mol. Errorbars are calculated as the SEM over 10 configurations. (b) Comparison of Δ*G_dil_*_−_*_to_*_−_*_dense_* between tyrosine (Tyr), neutral phosphotyrosine (pTyr^0^), charged phosphotyrosine (pTyr^-2^), lysine (Lys) and neutral lysine (Lys^0^). Errorbars are calculated as the SEM over 10 configurations. Δ*G_dil_*_−_*_to_*_−_*_dense_* for tyrosine and lysine are identical to those shown in Fig. 2a.

Moving on to the comparison among cationic and anionic residues, we found that Δ*G_*dil*-*to*-*dense*_* for arginine is significantly more favorable than lysine (Fig. 6a), supporting widespread observations that arginine promotes phase separation more effectively than lysine^10,12,16^. In contrast, the negatively charged residues aspartic acid and glutamic acid have comparable Δ*G_*dil*-*to*-*dense*_* values (Fig. 6a), explaining their similar contributions to phase separation, e.g. in hnRPA1 protein^16^.

### Charge asymmetry extends to modified amino acids

The asymmetry observed among the charged amino acids further prompts the question of whether the observation originates solely from the sign of the charge or from a combination of the other physicochemical features of arginine and lysine versus aspartic acid and glutamic acid. To better understand the role of the negative charge, we considered a comparison between tyrosine and two variants of phosphotyrosine, neutral phosphotyrosine (pTyr^0^) and charged phosphotyrosine (pTyr^-2^) which has a net charge of −2. We find that the addition of the phosphoryl group to the oxygen atom on the tyrosine side chain leads to an approximately twofold increase in Δ*G_*dil*-*to*-*dense*_* (Fig. 6b). Comparing 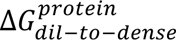 and 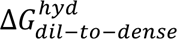 for tyrosine and neutral phosphotyrosine, we see that 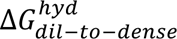 is more unfavorable for pTyr^0^ and can be predicted based on the fit of amino acid diameter (σ) and 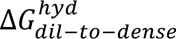 for all the neutral amino acids shown in Fig. 5d (Supplementary Fig. S5). However, this increase in the unfavorable 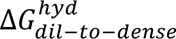 contribution is sufficiently offset by a more favorable 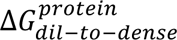 contribution (Supplementary Fig. S5), reflecting an enhancement in protein-analog interactions in the case of neutral phosphotyrosine as compared to tyrosine. Remarkably, the addition of the −2 charge to the phosphate group of the highly favorable neutral phosphotyrosine side chain analog, leading to pTyr^-2^, results in a 19.71 kJ/mol increase in Δ*G_*dil*-*to*-*dense*_* from −11.03 to +8.69 kJ/mol, indicating unfavorable transfer from the dilute phase to the condensate (Fig. 6b). As a control for the addition of positive charge, we compare the Δ*G_*dil*-*to*-*dense*_* of lysine and neutral lysine (Lys^0^). Removal of the positive charge from Lys still results in a marginally favorable Δ*G_*dil*-*to*-*dense*_* (Fig. 6b), but a comparison between lysine and neutral lysine reveals that addition of the positive charge leads to a decrease in Δ*G_*dil*-*to*-*dense*_* by 5.90 kJ/mol, underlining the preference for positive charge in the condensate microenvironment (Fig. 6b).

The findings of the different charge variants of unmodified and modified amino acids presented in this work clearly highlight that the observations made regarding the (un)favorability of transfer of the positive and negatively charged amino acids result in large part from the sign of the charge itself.

## Conclusions

In this work, we calculated the free energies associated with transferring all amino acid side chain analogs from the dilute phase to the dense phase of a model protein condensate, thereby providing a direct measure of residue-level driving forces underlying the formation of biomolecular condensates. We find that considering the dilute phase (as opposed to vacuum) as the reference state significantly improves the correlation between the calculated free energies and literature knowledge on residue-level contributions to phase separation. We find that within the condensate, the amino acids sample both solvent and protein rich environments (Fig. 7a), highlighting the role of solvent- and protein-mediated interactions in determining the (un)favorability of transfer. There is a delicate balance between a favorable protein contribution-stemming from protein interactions and the tendency of amino acids to leave the water-rich dilute phase-and an unfavorable hydration contribution, which arises from the energetic cost of forming cavities in the unique water structure inside the condensate (Fig. 7b). We observe that the hierarchy of amino acids in terms of their favorability in driving condensate formation is robust within the sequence space of polar-rich IDPs, but variations of about 2 kJ/mol between different protein sequences forming the condensate provide insight into the context-dependent nature of the sequence-level driving forces for phase separation. We additionally find that the highly favorable solvation of the negatively charged amino acids leads to the charge asymmetry in the contributions arising from the charged amino acids in their partitioning into the condensate. Positive charges favor the condensate microenvironment significantly more than negatively charged amino acids. The findings presented in this work will aid in the development of more robust physics-based theories to investigate sequence-dependent phase separation and will prompt further investigations into the potential role of charge as a modulator of biomolecular condensate formation in physiological contexts. These insights enhance our fundamental understanding of biomolecular condensate formation and could inform future experimental and computational studies aimed at manipulating phase separation processes in biological systems.

**Figure 7.**
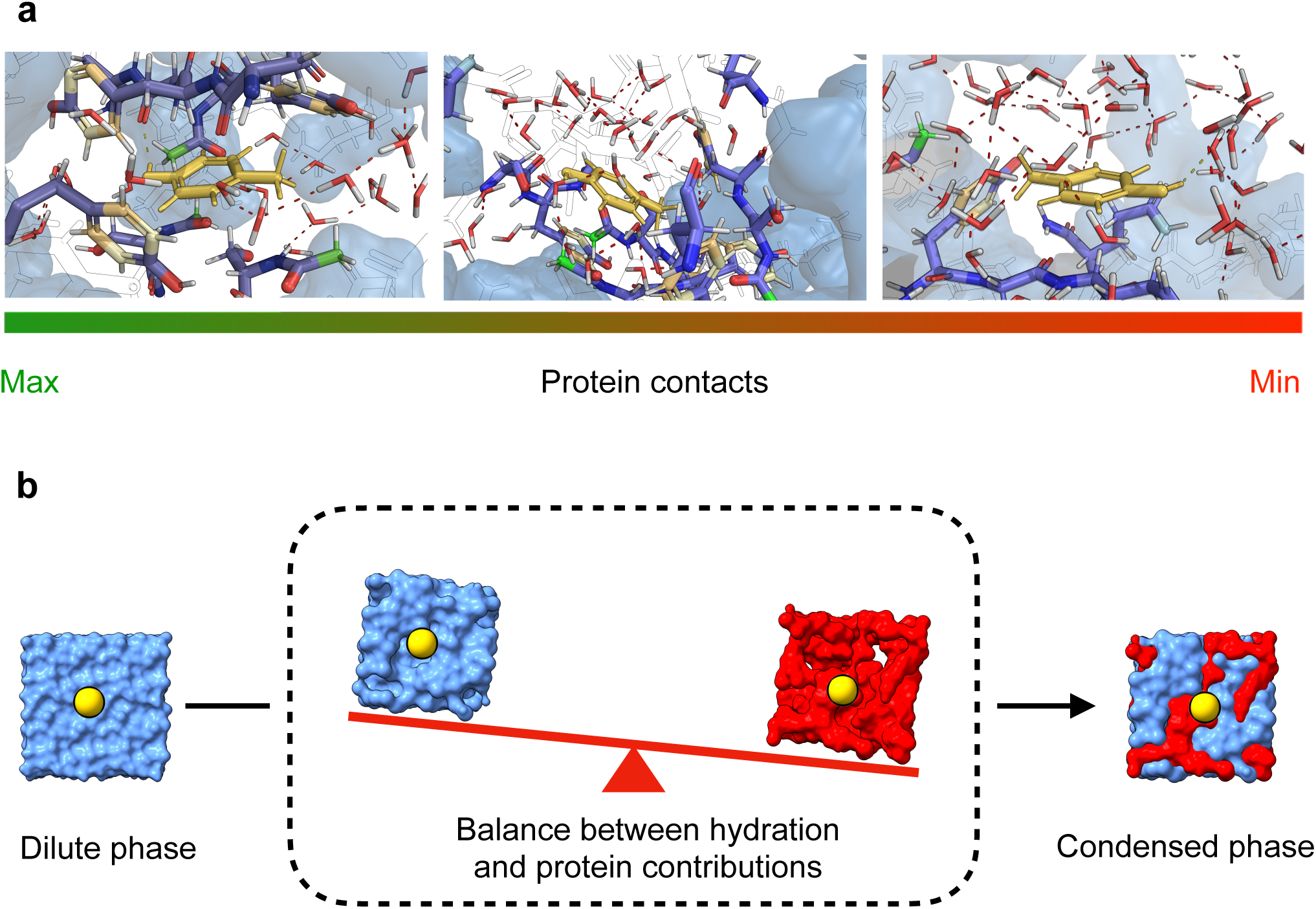
Complex interior of biomolecular condensates dictates residue-level contributions to phase separation. (a) Representative snapshots of the tyrosine side chain analog in SYGQ condensate as a function of protein-analog contacts. Red dashed lines show water-water hydrogen bonds while yellow dashed lines show analog-water hydrogen bonds. Protein atoms with distance from analog greater than 6Å are shown in black and white while water molecules with distance greater than 6Å from the analog are hidden for clarity. (b) Schematic highlighting the balance between unfavorable hydration and favorable protein mediated contributions in determining the net of amino acids from the dilute to the dense phase of biomolecular condensates.

## Methods

### Preparation of *in-silico* condensates

A single linear chain of the peptide sequence with ACE and NH_2_ capping groups at the N and C terminal ends was generated and saved as a PDB file using the tleap module in the AMBERTOOLS^75^ package. Following this, the based on the box size, the number of protein and water molecules were calculated based on the experimental measures of condensate density of the FUS-LC protein. The mass density of the protein was calculated ignoring the capping groups. Next, PACKMOL^76^ was used to insert the desired number of protein and water molecules into the simulation box. The resulting configuration is then minimized using GROMACS-2021.5^77^ steepest descent minimization, followed by 100 ps NVT and NPT runs with the v-rescale thermostat with set point 298 K for NVT, and v-rescale thermostat and the Berendsen barostat^78^ with set points 298 K and 1bar respectively. The equilibrated configurations are then run in the NPT ensemble using OPENMM-7.5^79^ for a runtime of 1 *μs* to ensure equilibration of the condensed phase prior to free energy calculations. The long runs on OPENMM-7.5^79^ are run with the AMBER03ws^46^ force-field in combination with TIP4P/2005^80^ water. Hydrogen masses are repartitioned to allow for a time step of 4fs. Short range non-bonded interactions are cut-off at 0.9 nm and long-range electrostatic interactions are treated with the PME method. Temperature is maintained using the Langevin integrator with time constant 1 ps and set point 298 K. Pressure is maintained at 1 bar using the isotropic Monte-Carlo Barostat implemented in OPENMM-7.5. Upon completion of the 1 *μs* runs, the final frame of the simulation is extracted for insertion of amino-acid analogs.

The condensate composed of the resilin-like-polypeptide (RLP) repeat unit (GRGDSPYS) chains is prepared following the same steps. However, due to the length of the sequence being double that of the SYGQ peptide, the number of chains required to achieve the same mass density of protein is half of that required for the SYGQ peptide. Upon preparing the system, minimizing and equilibrating, we run a 1 *μs* NPT simulation and see that the protein and water densities are similar to that of the SYGQ condensate (Supplementary Fig. S6)

### Generation and parametrization of amino acid side chain analogs

For the amino-acid side chain analogs, we began by building the residue of interest using the structure building tool in UCSF-Chimera^81^. The backbone atoms of the residue are deleted and then a PDB file of the side-chain atoms is saved. For the forcefield files, we follow the procedure detailed by Shirts et.al^82^. Namely, we take the original forcefield parameters (AMBER03ws) of the amino acid for which we are generating the analog forcefield files, delete the backbone atoms, and then increase or decrease the partial charges on the CB atom such that the sum of all the partial charges is equal to 0 for net neutral amino acids, +1 for cationic amino acids and −1 for anionic amino acids. These side chain PDB files are then inserted into the frame extracted from the condensate trajectory to get the configurations of the analog inserted within the condensates. For post translationally modified analogs (charged and neutral phosphotyrosine), the analogs were prepared using the same steps and modeled using the Forcefield_PTM forcefield^83^.

### Alchemical free energy calculations and analysis

Alchemical Free Energy calculations were done using GROMACS-2021.5. Electrostatic interactions were decoupled first, followed by vdw interactions. The total number of windows used for decoupling of the analog from the environment was 16. The exact values for electrostatic decoupling were 0.00, 0.20, 0.40, 0.60, 0.80, 1.00, 1.00, 1.00, 1.00, 1.00, 1.00, 1.00, 1.00, 1.00, 1.00, 1.00 and for vdw decoupling were 0.00, 0.00, 0.00, 0.00, 0.00, 0.00, 0.10, 0.20, 0.30, 0.40, 0.50, 0.60, 0.70, 0.80, 0.90, 1.00. Decoupling simulations were run for 5ns for each replica or configuration using the Langevin integrator with temperature set to 298 K and time constant 1 ps. Pressure was maintained at 1 bar using the Parinello-Rahman barostat with a coupling constant of 1 ps. Soft-core potentials were used for LJ interactions to avoid singularities with alpha parameter set to 0.4. For all analogs apart from Ala, hydrogen mass repartitioning was done, and all bonds were constrained using LINCS allowing for a time step of 5fs (2fs for Ala). Short range interactions were treated using a cutoff of 1.2 nm, while long range electrostatic interactions were treated using PME^84^ with a real space cutoff of 1.2 nm. Free energies were calculated using the MBAR^85^ estimator in the alchemlyb python package.

### Calculation of *ΔG_vac-to-dil_*

A cubic box of side 3 nm was filled with water molecules then equilibrated in OPENMM-7.5 with the same run parameters as for the equilibration of the *in silico* condensate. Following this equilibration of the water box, each of the amino acids were randomly inserted within the box and the system was minimized then equilibrated in the NPT ensemble using the Parinello-Rahman barostat with coupling constant of 1 ps for pressure control and the Langevin integrator with coupling constant 1 ps for temperature control. Following the equilibration the alchemical free energy calculations were done with the run parameters detailed above. Three such replicas were run for each of the amino acid side chain analogs. Δ*G_*vac*-*to*-*dil*_* values reported in the article are the average of these three replicas with the uncertainties reported as the standard error of the mean across the three replicas.

### Assessment of the impact of local protein density variations on calculation of *ΔG_*vac*-*to*-*dense*_*

The observation of local variations in protein and water densities within the condensate system demonstrates that, on the timescales accessible to current atomic-resolution simulations for free energy calculations, BCs do not behave as bulk-like media. This heterogeneity suggests that the calculated Δ*G_*vac*-*to*-*dense*_* values will depend on the local environment sampled by the side chain analog within the simulation. To investigate this dependence, we focused on the side chain analog of tryptophan due to its large size. We calculated Δ*G_*vac*-*to*-*dense*_* as a function of simulation time in each alchemical window while maintaining the same starting configuration of the analog inserted within the condensate. We found that after 5 ns of simulation time per window, estimates of Δ*G_*vac*-*to*-*dense*_* showed no appreciable difference even up to 100 ns of simulation time per window indicating convergence of the free energy calculations within 5 ns. (Supplementary Fig. S7). Next, we calculated Δ*G_*vac*-*to*-*dense*_* for three different initial positions of the tryptophan analog within the condensate, using a fixed simulation time of 5 ns per window. We observed that variations in Δ*G_*vac*-*to*-*dense*_* as a function of tryptophan position within the condensed phase were on the order of 3 kJ/mol (Supplementary Fig. S7) indicating a dependence of Δ*G_*vac*-*to*-*dense*_* on the local environment of the analog within the condensate. To obtain reliable estimates of Δ*G_*vac*-*to*-*dense*_*, and account for this position dependence, we devised a sampling protocol where each analog is inserted into the condensate, and the system is then equilibrated for 1 *μs* in the NPT ensemble using OPENMM-7.5 using the same run parameters as in the simulations to equilibrate of the condensate to allow diffusion of the analog throughout the condensate. Upon completion of the run, the number of contacts between the analog and protein are calculated. The number of analog-protein contacts are computed based on a distance criterion such that any heavy atom pair from the analog and protein respectively that are within 6Å of each other are counted as a contact. A histogram of 10 bins is then generated based on the number of analog-protein contacts observed during the trajectory (Supplementary Fig. S8). From each bin of the resulting histogram a frame is selected which serves as a starting configuration for the alchemical free energy calculations, ensuring coverage of environments ranging from protein-rich to protein-depleted regions within the condensate. We then conducted alchemical free energy calculations for each of these configurations, using a simulation time of 5 ns per window, to compute Δ*G_*vac*-*to*-*dense*_*. This was done for both the SYGQ and RLP condensates. Δ*G_*vac*-*to*-*dense*_* values reported are the mean of the Δ*G_*vac*-*to*-*dense*_* values calculated from the 10 different starting configurations and uncertainties reported are the standard error of the mean across the 10 different configurations for each of the amino acid side chain analogs.

### Calculation of 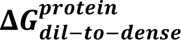

To estimate the transfer from vacuum to bulk protein, water molecules were deleted from the final frame of the 1 *μs* simulation of the condensate. Water molecules were removed and then the analog was inserted. Spherical flat bottom restraints with radius 1 nm and force constant 1000 kJ/mol were then applied to all non-hydrogen atoms of the peptide chains to prevent collapse of the chains which would lead to higher protein densities. The resulting analog in position retrained peptide configuration was then equilibrated using the Langevin Integrator in GROMACS at 298K with friction 1 ps for 2.5 ns to allow for movement of the analog. The equilibrated configuration was then used for the alchemical free energy calculations. To prevent collapse of the peptide chains, the simulations to estimate free energy were done with no pressure coupling. Apart from the pressure coupling, all run parameters, and the analysis protocol detailed in the alchemical free energy calculations and analysis section was followed. Uncertainty estimates on free energy were made from 5 insertions run.

### Calculation of interfacial electrostatic potential and Galvani potentials

A surface or interfacial correction is required in alchemical free energy calcualtions to account for the absence of an interface between the two phases being considered. This interfacial contribution can be written in terms of the electrostatic potential drop going from one phase to the other,

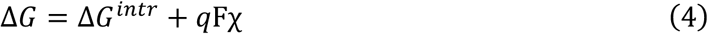

Where Δ*G*^*iintr*^ is the free energy measured in the simulation, q is the charge of the amino acid, F is Faraday’s constant and *χ* is the electrostatic potential difference between the two phases. The factor *Fχ* is the Galvani potential between the two phases (Φ_*G*_). For neutral amino acids, since q=0, Δ*G* = Δ*G*^*iintr*^. However, for charged amino acids, based on the sign of the charge, the free energies must be corrected. For the charged amino acids, we can write this corrected transfer free energy from the dilute to the dense phase as,

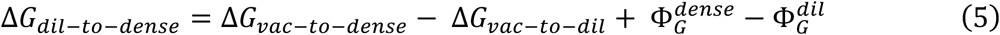

Where 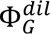 and 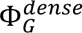 are the Galvani potential differences across the vacuum-to-water and vacuum-to-condensate phases respectively in the vacuum-to-dilute and vacuum-to-dense direction respectively. To calculate the Galvani potential of the dilute phase, we first compute the electrostatic potential (*χ*) difference across the vacuum-to-water interface using the approach detailed in Zhang et.al^42^. Namely, we consider equilibrate a 5 nm box of pure solvent and a 5 nm box of the condensate system for the dilute and dense phase in OPENMM-7.5 using the same run parameters as detailed in the equilibration of the *in silico* condensate. Following this, we extract the last frame of the equilibration trajectory and extend the box in the z dimension to 15 nm leading to a cuboidal simulation cell of 5×5×15 nm dimension. Then the systems are run in the NVT ensemble on GROMACS for 10 ns using the Langevin integrator with time constant 1 ps. From this trajectory, we compute the electrostatic potential of the dilute and dense phases over the last 5 ns of simulation time using the command gmx potential with the -symm flag to symmetrize the profiles. We estimate *χ*^*dil*^ and *χ*^*dense*^ as −0.498 and −0.285 V and using these values (Supplementary Fig. S9), we get 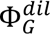 and 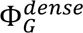 equal to −48.06 and −27.55 kJ/mol/e in the vacuum-to-dilute and vacuum-to-condensate direction respectively. When we use the calculated 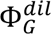 to correct the intrinsic Δ*G_*vac*-*to*-*dil*_* values for the charged residues, we observe a good match with previous estimates of solvation free energies of the charged amino acids (Supplementary Fig. S10). Substituting the values of 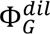 and 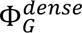 in equation 5 along with the charges, we get the corrected free energies for the charged amino acids.

## Acknowledgements

This work was supported by National Institute of General Medical Science of the National Institute of Health award R35GM153388. We also acknowledge the support provided by the Welch Foundation (A-2113-202203311). Texas A&M High Performance Research Computing provided computational resources used in this work. We acknowledge Dr. Young C. Kim (Naval Research Lab), Dr. Benjamin Schuster (Rutgers University), Dr. Nicolas Fawzi (Brown University), Dr. Priyesh Mohanty (Texas A&M University), Dr. Dinesh Sundaravadivelu Devarajan (Texas A&M University), and Azamat Rizuan (Texas A&M University) for providing feedback on the earlier version of this manuscript. We also gratefully acknowledge Dr. Priyesh Mohanty (Texas A&M University) for providing the forcefield parameters for the amino acid side chain analogs.

## References

1 Boeynaems, S. et al. Protein phase separation: a new phase in cell biology. Trends in cell biology 28, 420–435 (2018).

2 Banani, S. F., Lee, H. O., Hyman, A. A. & Rosen, M. K. Biomolecular condensates: organizers of cellular biochemistry. Nature reviews Molecular cell biology 18, 285–298 (2017).

3 Patel, A. et al. A liquid-to-solid phase transition of the ALS protein FUS accelerated by disease mutation. Cell 162, 1066–1077 (2015).

4 Conicella, A. E., Zerze, G. H., Mittal, J. & Fawzi, N. L. ALS mutations disrupt phase separation mediated by α-helical structure in the TDP-43 low-complexity C-terminal domain. Structure 24, 1537–1549 (2016).

5 Mackenzie, I. R. et al. TIA1 mutations in amyotrophic lateral sclerosis and frontotemporal dementia promote phase separation and alter stress granule dynamics. Neuron 95, 808–816. e809 (2017).

6 Shin, Y. & Brangwynne, C. P. Liquid phase condensation in cell physiology and disease. Science 357, eaaf4382 (2017).

7 Patkar, S. S., Garcia Garcia, C., Palmese, L. L. & Kiick, K. L. Sequence-Encoded Differences in Phase Separation Enable Formation of Resilin-like Polypeptide-Based Microstructured Hydrogels. Biomacromolecules 24, 3729–3741 (2023).

8 Varanko, A., Saha, S. & Chilkoti, A. Recent trends in protein and peptide-based biomaterials for advanced drug delivery. Advanced drug delivery reviews 156, 133–187 (2020).

9 Liu, A. P. et al. The living interface between synthetic biology and biomaterial design. Nature materials 21, 390–397 (2022).

10 Wang, J. et al. A molecular grammar governing the driving forces for phase separation of prion-like RNA binding proteins. Cell 174, 688–699. e616 (2018).

11 Nott, T. J. et al. Phase transition of a disordered nuage protein generates environmentally responsive membraneless organelles. Molecular cell 57, 936–947 (2015).

12 Schuster, B. S. et al. Identifying sequence perturbations to an intrinsically disordered protein that determine its phase-separation behavior. Proceedings of the National Academy of Sciences 117, 11421–11431 (2020).

13 Rekhi, S. et al. Expanding the molecular language of protein liquid-liquid phase separation. Nature Chemistry (2024). 10.1038/s41557-024-01489-x

14 Quiroz, F. G. & Chilkoti, A. Sequence heuristics to encode phase behaviour in intrinsically disordered protein polymers. Nature materials 14, 1164–1171 (2015).

15 Dzuricky, M., Rogers, B. A., Shahid, A., Cremer, P. S. & Chilkoti, A. De novo engineering of intracellular condensates using artificial disordered proteins. Nature chemistry 12, 814–825 (2020).

16 Bremer, A. et al. Deciphering how naturally occurring sequence features impact the phase behaviours of disordered prion-like domains. Nature Chemistry 14, 196–207 (2022).

17 Dignon, G. L., Zheng, W., Kim, Y. C., Best, R. B. & Mittal, J. Sequence determinants of protein phase behavior from a coarse-grained model. PLoS computational biology 14, e1005941 (2018).

18 Joseph, J. A. et al. Physics-driven coarse-grained model for biomolecular phase separation with near-quantitative accuracy. Nature Computational Science 1, 732–743 (2021).

19 Regy, R. M., Thompson, J., Kim, Y. C. & Mittal, J. Improved coarse-grained model for studying sequence dependent phase separation of disordered proteins. Protein Science 30, 1371–1379 (2021).

20 Tesei, G., Schulze, T. K., Crehuet, R. & Lindorff-Larsen, K. Accurate model of liquid– liquid phase behavior of intrinsically disordered proteins from optimization of single-chain properties. Proceedings of the National Academy of Sciences 118, e2111696118 (2021).

21 Houston, L., Phillips, M., Torres, A., Gaalswyk, K. & Ghosh, K. Physics-Based Machine Learning Trains Hamiltonians and Decodes the Sequence–Conformation Relation in the Disordered Proteome. Journal of Chemical Theory and Computation (2024).

22 Dekker, M., Van der Giessen, E. & Onck, P. R. Phase separation of intrinsically disordered FG-Nups is driven by highly dynamic FG motifs. Proceedings of the National Academy of Sciences 120, e2221804120 (2023).

23 Das, S., Lin, Y.-H., Vernon, R. M., Forman-Kay, J. D. & Chan, H. S. Comparative roles of charge, π, and hydrophobic interactions in sequence-dependent phase separation of intrinsically disordered proteins. Proceedings of the National Academy of Sciences 117, 28795–28805 (2020).

24 Tesei, G. & Lindorff-Larsen, K. Improved predictions of phase behaviour of intrinsically disordered proteins by tuning the interaction range. Open Research Europe 2 (2022).

25 Betancourt, M. R. & Thirumalai, D. Pair potentials for protein folding: choice of reference states and sensitivity of predicted native states to variations in the interaction schemes. Protein science 8, 361–369 (1999).

26 Lau, K. F. & Dill, K. A. A lattice statistical mechanics model of the conformational and sequence spaces of proteins. Macromolecules 22, 3986–3997 (1989).

27 Miyazawa, S. & Jernigan, R. L. An empirical energy potential with a reference state for protein fold and sequence recognition. Proteins: Structure, Function, and Bioinformatics 36, 357–369 (1999).

28 Wolfenden, R., Andersson, L., Cullis, P. & Southgate, C. Affinities of amino acid side chains for solvent water. Biochemistry 20, 849–855 (1981).

29 Radzicka, A. & Wolfenden, R. Comparing the polarities of the amino acids: side-chain distribution coefficients between the vapor phase, cyclohexane, 1-octanol, and neutral aqueous solution. Biochemistry 27, 1664-1670 (1988).

30 Rekhi, S. et al. Role of strong localized vs weak distributed interactions in disordered protein phase separation. The Journal of Physical Chemistry B 127, 3829–3838 (2023).

31 Statt, A., Casademunt, H., Brangwynne, C. P. & Panagiotopoulos, A. Z. Model for disordered proteins with strongly sequence-dependent liquid phase behavior. The Journal of chemical physics 152 (2020).

32 Michels, J. J., Brzezinski, M., Scheidt, T., Lemke, E. A. & Parekh, S. H. Role of solvent compatibility in the phase behavior of binary solutions of weakly associating multivalent polymers. Biomacromolecules 23, 349–364 (2021).

33 Pesce, F. et al. Design of intrinsically disordered protein variants with diverse structural properties. Science Advances 10, eadm9926 (2024).

34 Devarajan, D. S. et al. Sequence-dependent material properties of biomolecular condensates and their relation to dilute phase conformations. Nature Communications 15 (2024).

35 Lin, Y., Currie, S. L. & Rosen, M. K. Intrinsically disordered sequences enable modulation of protein phase separation through distributed tyrosine motifs. Journal of Biological Chemistry 292, 19110–19120 (2017).

36 Dannenhoffer-Lafage, T. & Best, R. B. A data-driven hydrophobicity scale for predicting liquid–liquid phase separation of proteins. The Journal of Physical Chemistry B 125, 4046–4056 (2021).

37 Latham, A. P. & Zhang, B. Consistent force field captures homologue-resolved HP1 phase separation. Journal of chemical theory and computation 17, 3134–3144 (2021).

38 Urry, D. W. et al. Hydrophobicity scale for proteins based on inverse temperature transitions. Biopolymers: Original Research on Biomolecules 32, 1243–1250 (1992).

39. Boeynaems, S., et al. Aberrant phase separation is a common killing strategy of positively charged peptides in biology and human disease. bioRxiv (2023).

40 Mukherjee, S. & Schäfer, L. V. Thermodynamic forces from protein and water govern condensate formation of an intrinsically disordered protein domain. Nature Communications 14, 5892 (2023).

41 Zhang, H. et al. Free-energy calculations of ionic hydration consistent with the experimental hydration free energy of the proton. The Journal of Physical Chemistry Letters 8, 2705–2712 (2017).

42 Zhang, H., Yin, C., Jiang, Y. & van der Spoel, D. Force field benchmark of amino acids: I. hydration and diffusion in different water models. Journal of chemical information and modeling 58, 1037–1052 (2018).

43 Leung, K. Surface potential at the air− water interface computed using density functional theory. The Journal of Physical Chemistry Letters 1, 496–499 (2010).

44 Lin, Y.-L., Aleksandrov, A., Simonson, T. & Roux, B. An overview of electrostatic free energy computations for solutions and proteins. Journal of chemical theory and computation 10, 2690–2709 (2014).

45 Remsing, R. C., Baer, M. D., Schenter, G. K., Mundy, C. J. & Weeks, J. D. The role of broken symmetry in solvation of a spherical cavity in classical and quantum water models. The journal of physical chemistry letters 5, 2767–2774 (2014).

46 Best, R. B., Zheng, W. & Mittal, J. Balanced protein–water interactions improve properties of disordered proteins and non-specific protein association. Journal of chemical theory and computation 10, 5113–5124 (2014).

47 Paloni, M., Bailly, R., Ciandrini, L. & Barducci, A. Unraveling molecular interactions in liquid–liquid phase separation of disordered proteins by atomistic simulations. The Journal of Physical Chemistry B 124, 9009–9016 (2020).

48 Workman, R. J. & Pettitt, B. M. Thermodynamic Compensation in Peptides Following Liquid–Liquid Phase Separation. The Journal of Physical Chemistry B 125, 6431–6439 (2021).

49 Rauscher, S. & Pomès, R. The liquid structure of elastin. Elife 6, e26526 (2017).

50 Martin, E. W. & Mittag, T. Relationship of sequence and phase separation in protein low-complexity regions. Biochemistry 57, 2478–2487 (2018).

51 Murthy, A. C. et al. Molecular interactions underlying liquid− liquid phase separation of the FUS low-complexity domain. Nature structural & molecular biology 26, 637–648 (2019).

52 Zheng, W. et al. Molecular details of protein condensates probed by microsecond long atomistic simulations. The Journal of Physical Chemistry B 124, 11671–11679 (2020).

53 Galvanetto, N. et al. Extreme dynamics in a biomolecular condensate. Nature 619, 876–883 (2023).

54 Dar, F. et al. Biomolecular condensates form spatially inhomogeneous network fluids. Nature communications 15, 3413 (2024).

55 Johnson, C. N. et al. Insights into Molecular Diversity within the FUS/EWS/TAF15 Protein Family: Unraveling Phase Separation of the N-Terminal Low-Complexity Domain from RNA-Binding Protein EWS. Journal of the American Chemical Society

56 Sabari, B. R. et al. Coactivator condensation at super-enhancers links phase separation and gene control. Science 361, eaar3958 (2018).

57 Mohanty, P. et al. A synergy between site-specific and transient interactions drives the phase separation of a disordered, low-complexity domain. Proceedings of the National Academy of Sciences 120, e2305625120 (2023).

58. Rizuan, A., et al. Structural details of helix-mediated TDP-43 C-terminal domain multimerization. bioRxiv (2024).

59. Wake, N., et al. Expanding the molecular grammar of polar residues and arginine in FUS prion-like domain phase separation and aggregation. bioRxiv (2024).

60 Martin, E. W. et al. Valence and patterning of aromatic residues determine the phase behavior of prion-like domains. Science 367, 694–699 (2020).

61 Ambadi Thody, S., et al. Small-molecule properties define partitioning into biomolecular condensates. Nature Chemistry, 1–9 (2024).

62 Kilgore, H. R. et al. Distinct chemical environments in biomolecular condensates. Nature Chemical Biology 20, 291–301 (2024).

63 Kim, Y. C. & Hummer, G. Coarse-grained models for simulations of multiprotein complexes: application to ubiquitin binding. Journal of molecular biology 375, 1416–1433 (2008).

64 De Sancho, D. & Lopez, X. Crossover in Aromatic Amino Acid Interaction Strength: Tyrosine vs. Phenylalanine in Biomolecular Condensates. bioRxiv, 2024.2010. 2018.619001 (2024).

65 Errington, J. R. & Debenedetti, P. G. Relationship between structural order and the anomalies of liquid water. Nature 409, 318–321 (2001).

66 Rasaiah, J. C., Garde, S. & Hummer, G. Water in nonpolar confinement: From nanotubes to proteins and beyond. Annu. Rev. Phys. Chem. 59, 713–740 (2008).

67 Rego, N. B. & Patel, A. J. Understanding hydrophobic effects: Insights from water density fluctuations. Annual Review of Condensed Matter Physics 13, 303–324 (2022).

68 Jamadagni, S. N., Godawat, R. & Garde, S. Hydrophobicity of proteins and interfaces: Insights from density fluctuations. Annual review of chemical and biomolecular engineering 2, 147–171 (2011).

69 Chandler, D. Interfaces and the driving force of hydrophobic assembly. Nature 437, 640–647 (2005).

70 Ashbaugh, H. S., Truskett, T. M. & Debenedetti, P. G. A simple molecular thermodynamic theory of hydrophobic hydration. The Journal of chemical physics 116, 2907–2921 (2002).

71 Hummer, G., Pratt, L. R. & Garcia, A. E. Free energy of ionic hydration. The Journal of Physical Chemistry 100, 1206–1215 (1996).

72 Baruch Leshem, A., et al. Biomolecular condensates formed by designer minimalistic peptides. Nature Communications 14, 421 (2023).

73. Crabtree, M. D., et al. Ion binding with charge inversion combined with screening modulates DEAD box helicase phase transitions. Cell Reports 42 (2023).

74 Saar, K. L. et al. Learning the molecular grammar of protein condensates from sequence determinants and embeddings. Proceedings of the National Academy of Sciences 118, e2019053118 (2021).

75 D.A. Case, H. M. A., K. Belfon, I.Y. Ben-Shalom, S.R. Brozell, D.S. Cerutti, T.E. Cheatham, III, G.A. et al. Amber 2021. University of California, San Francisco (2021).

76 Martínez, L., Andrade, R., Birgin, E. G. & Martínez, J. M. PACKMOL: A package for building initial configurations for molecular dynamics simulations. Journal of computational chemistry 30, 2157–2164 (2009).

77 Abraham, M. J. et al. GROMACS: High performance molecular simulations through multi-level parallelism from laptops to supercomputers. SoftwareX 1, 19–25 (2015).

78 Berendsen, H. J., Postma, J. v., Van Gunsteren, W. F., DiNola, A. & Haak, J. R. Molecular dynamics with coupling to an external bath. The Journal of chemical physics 81, 3684–3690 (1984).

79 Eastman, P. et al. OpenMM 7: Rapid development of high performance algorithms for molecular dynamics. PLoS computational biology 13, e1005659 (2017).

80 Abascal, J. L. & Vega, C. A general purpose model for the condensed phases of water: TIP4P/2005. The Journal of chemical physics 123, 234505 (2005).

81 Pettersen, E. F. et al. UCSF Chimera—a visualization system for exploratory research and analysis. Journal of computational chemistry 25, 1605–1612 (2004).

82 Shirts, M. R., Pitera, J. W., Swope, W. C. & Pande, V. S. Extremely precise free energy calculations of amino acid side chain analogs: Comparison of common molecular mechanics force fields for proteins. The Journal of chemical physics 119, 5740–5761 (2003).

83 Khoury, G. A., Thompson, J. P., Smadbeck, J., Kieslich, C. A. & Floudas, C. A. Forcefield_PTM: Ab initio charge and AMBER forcefield parameters for frequently occurring post-translational modifications. Journal of chemical theory and computation 9, 5653–5674 (2013).

84 Darden, T., York, D. & Pedersen, L. Particle mesh Ewald: An N⋅ log (N) method for Ewald sums in large systems. The Journal of chemical physics 98, 10089–10092 (1993).

85 Shirts, M. R. & Chodera, J. D. Statistically optimal analysis of samples from multiple equilibrium states. The Journal of chemical physics 129 (2008).

