## Supplementary Information for "Amino Acid Transfer Free Energies Reveal Thermodynamic Driving Forces in Biomolecular Condensate Formation"

### Supporting Figures

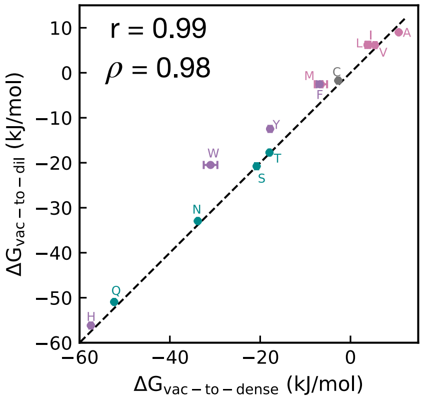


***Figure S1.*** *Correlation between* $\Delta G_{vac-to-dense}$ *and* $\Delta G_{vac-to-dil}$ *for all neutral amino acids. Pearson correlation coefficient (r) and Spearman rank correlation coefficient (ρ) are shown as measures of correlation.*

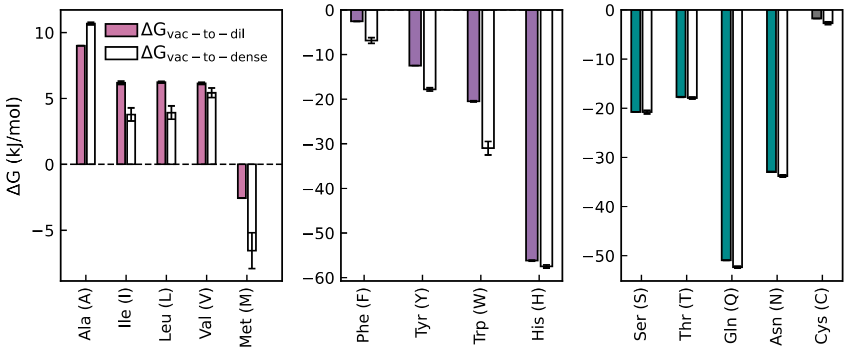


***Figure S2.*** *Comparison between* $\Delta G_{vac-to-dil}$ *and* $\Delta G_{vac-to-dense}$ *for aliphatic, aromatic, polar and other amino acids.* $\Delta G_{vac-to-dil}$ *values are shown as filled bars, while* $\Delta G_{vac-to-dense}$ *values are shown as empty bars.*

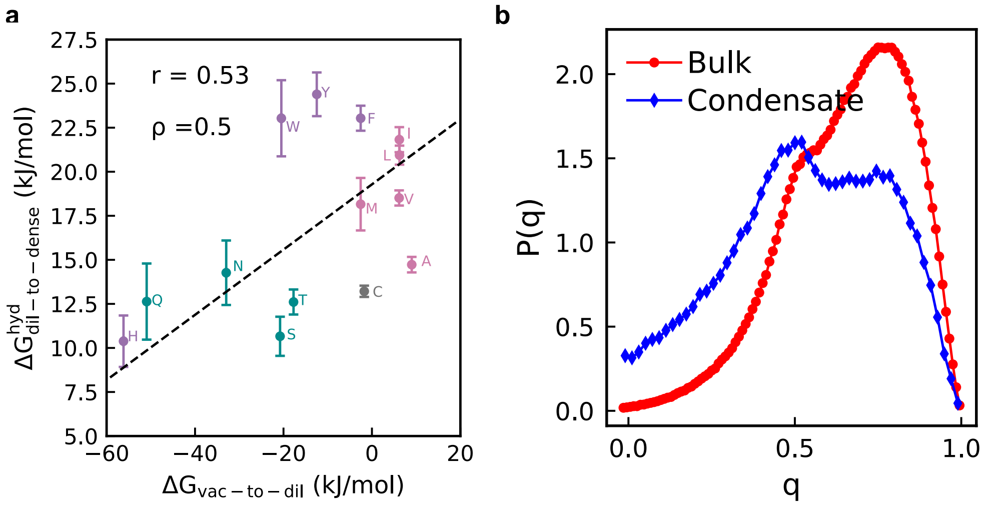


***Figure S3.*** *(a) Correlation between* $\Delta G^{hyd}_{dil-to-dense}$ *and* $\Delta G_{vac-to-dil}$ *for all neutral amino acids. Pearson correlation coefficient (r) and Spearman rank correlation coefficient* ($\rho$) *are shown as measures of correlation. (b) Comparison of probability density distributions of the tetrahedral order parameter for water in a pure water (bulk) system and within the condensate. A q value of 1 indicates a perfect tetrahedral arrangement, while 0 indicates ideal gas-like behavior.*

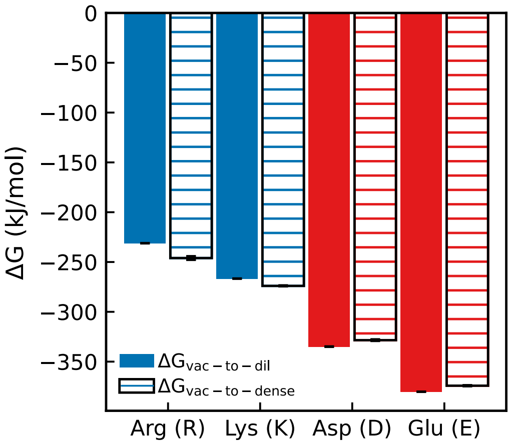


***Figure S4.*** *Comparison between* $\Delta G_{vac-to-dil}$ *(filled bars) and* $\Delta G_{vac-to-dense}$ *(hatched bars) in kJ/mol) for the charged amino acids. Errorbars are the SEM over 10 insertions for* $\Delta G_{vac-to-dense}$ *and over 3 insertions for* $\Delta G_{vac-to-dil}$.

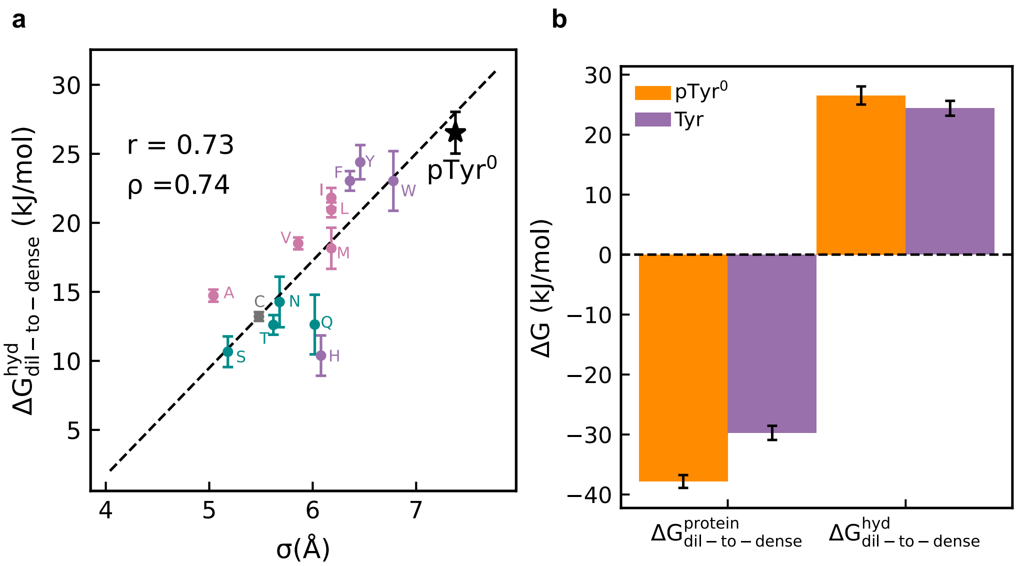


***Figure S5.*** *(a) Correlation between amino acid diameter and* $\Delta G^{hyd}_{dil-to-dense}$ *as shown in Fig. 3c of the main text. The black star shows neutral phosphotyrosine (pTyr⁰) (b) Comparison between* $\Delta G^{protein}_{dil-to-dense}$ *and* $\Delta G^{hyd}_{dil-to-dense}$ *of Tyr and neutral phosphotyrosine (pTyr⁰).*

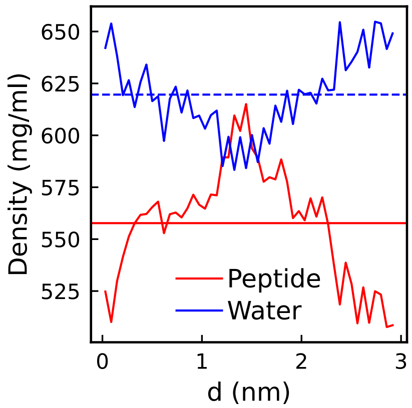


***Figure S6.*** *Average mass densities of peptide and water in a condensate composed of RLP repeat unit peptides. Densities are estimated from a 1µs NPT run. Dashed blue and solid red horizontal lines iindicate average densities of water and peptide respectively within the simulation cell.*

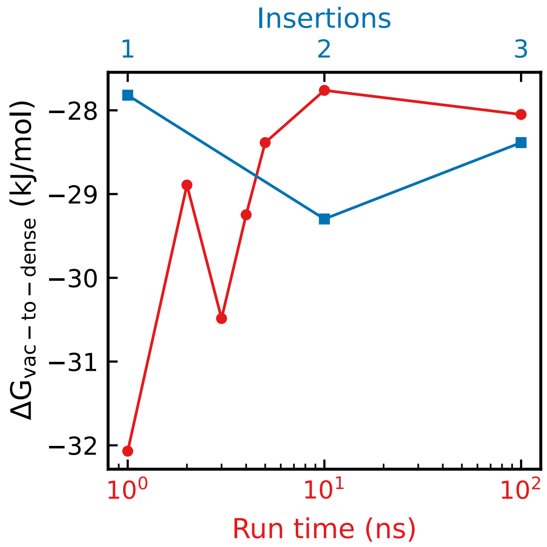


***Figure S7.*** *Variation of $\Delta G_{vac-to-dense}$ of the tryptophan side chain analog within the SYGQ peptide condensate as a function of run time (red) and insertion position (blue). Run times tested are 1,2,3,4,5,10 and 100 ns per sampling window.*

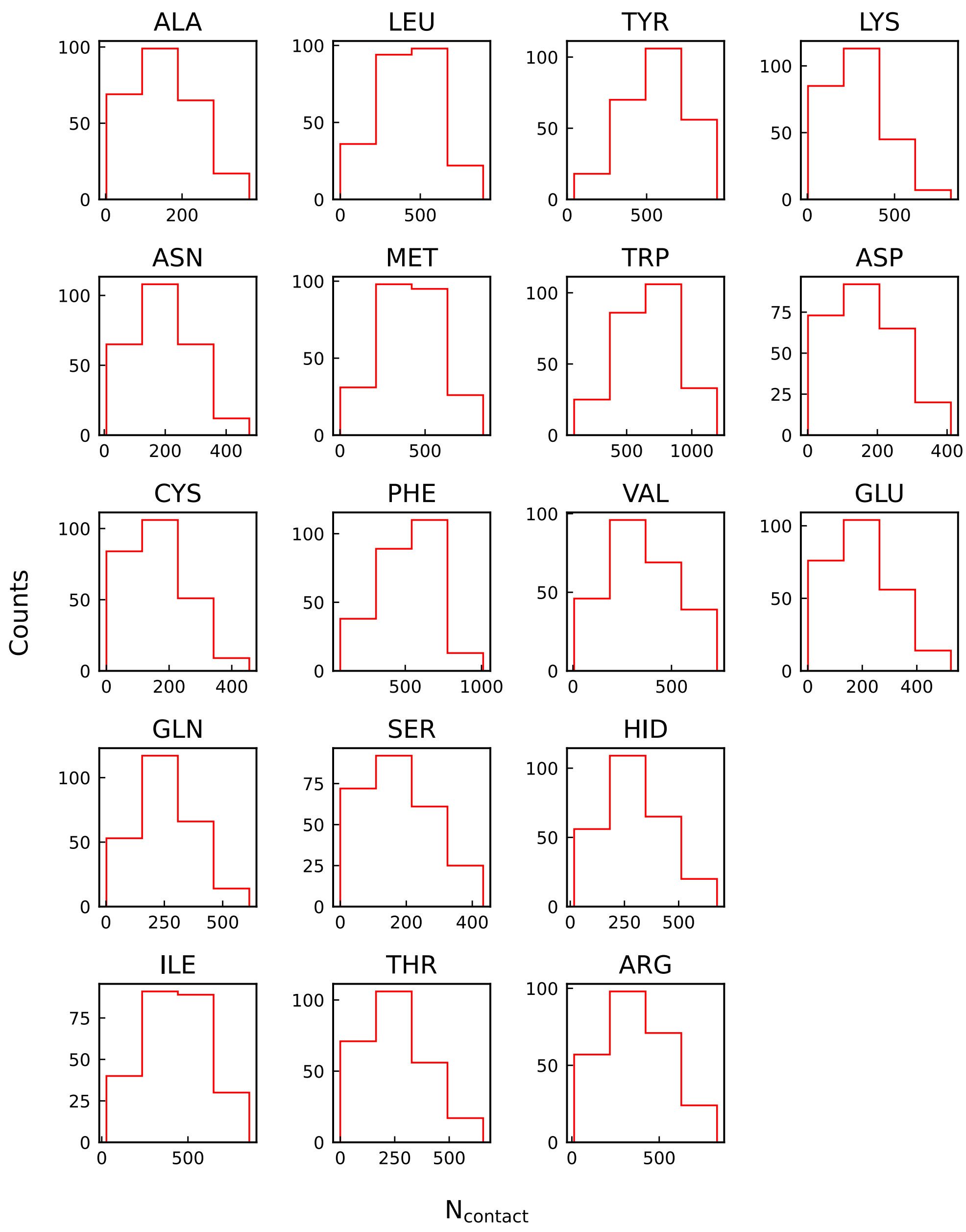


***Figure S8.*** *Histograms of number of analog-peptide contacts for all the amino acid side chain analogs in the SYGQ condensate. Histograms are constructed with 10 bins such that one configuration is selected from each bin as an initial condition for the estimation of* $\Delta G_{cond}$. *Contacts are calculated from a* $1\mu s$ *NPT simulation of the analog within the SYGQ condensate.*

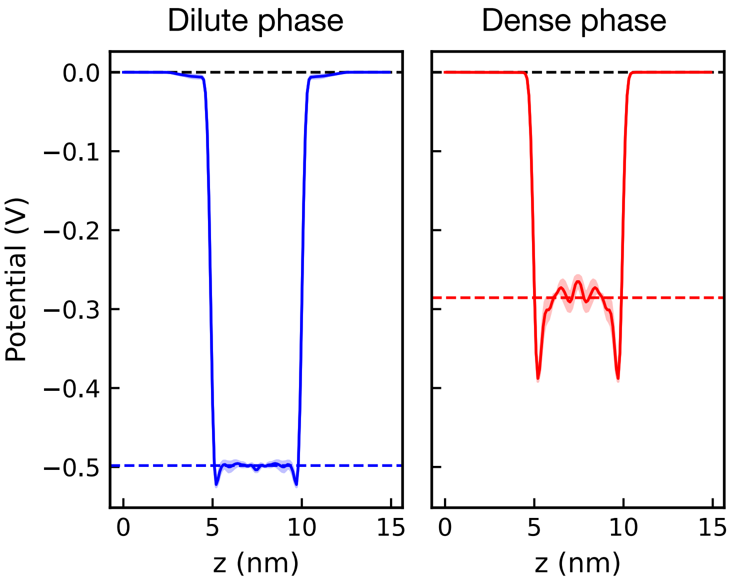


***Figure S9.*** *Electrostatic potential from the vacuum to the dilute (blue) and dense (red) phase along the z dimension of the simulation box. The vacuum electrostatic potential is set to 0. The dashed lines indicate the average value over 3 different simulations taken from the z dimension range of 5.5 to 9.5 nm.*

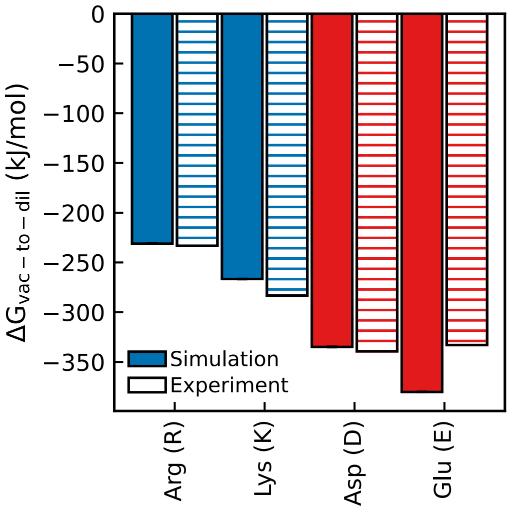


***Figure S10.*** *Comparison between* $\Delta G_{vac-to-dil}$ *for the charged amino acids between simulation (filled bars) and experiment (hatched bars) for the charged amino acids. Experimental values for the amino acids are taken from Fossat et.al[1] and Zhang et.al[2].*

### References


1 Fossat, M. J., Zeng, X. & Pappu, R. V. Uncovering differences in hydration free energies and structures for model compound mimics of charged side chains of amino acids. *The Journal of Physical Chemistry B* **125**, 4148-4161 (2021).

2 Zhang, H., Yin, C., Jiang, Y. & van der Spoel, D. Force field benchmark of amino acids: I. hydration and diffusion in different water models. *Journal of chemical information and modeling* **58**, 1037-1052 (2018).
